# High prevalence of CNS-directed autoantibodies in patients with schizophrenia

**DOI:** 10.64898/2026.05.04.722731

**Authors:** Katlyn Nemani, Jillian R. Jaycox, Ugur Akcan, Benjamin M. Schuman, Sung M. Yeon, Nina Harano, Kai Qin, Alison A. Notestine, Sean M. Carroll, Brent S. McKenzie, Leon Furchtgott, Adrienne C. Lahti, Richard W. Tsien, Dritan Agalliu, Donald C. Goff, Aaron M. Ring

**Affiliations:** Department of Psychiatry, New York University Langone Medical Center, New York, NY, USA; Nathan S. Kline Institute for Psychiatric Research, Orangeburg, NY, USA; Division of Translational Science and Therapeutics, Fred Hutchinson Cancer Center, Seattle, WA, USA; Department of Neurology, Columbia University Irving Medical Center, New York, NY, USA; Department of Neuroscience and Physiology, New York University Langone Medical Center, New York, USA; Seranova Bio, South San Francisco, CA, USA; Department of Psychiatry and Behavioral Neurobiology, School of Medicine, University of Alabama at Birmingham, Birmingham, AL, USA

## Abstract

Schizophrenia is a severe neuropsychiatric disorder whose etiology and biological heterogeneity remain poorly understood. Immune dysregulation has long been implicated, but the breadth and clinical significance of autoantibody responses remain unclear beyond rare individual examples. Here we use Rapid Extracellular Antigen Profiling–a proteome-scale assay for autoantibodies against extracellular and secreted proteins–to profile 352 individuals with schizophrenia and 971 community controls. We find that schizophrenia is marked by a striking elevation in extracellular autoantibody burden, present near disease onset, and approaching nearly twice control levels in the most severely ill patients. This burden increases with age in a pattern that diverges from controls and targets central nervous system antigens, including neuroactive receptors, neuronal ion channels, proteins involved in synaptic function, and blood–brain barrier integrity. Autoantibodies against blood–brain barrier antigens impair barrier function *ex vivo* and are associated with broader central nervous system autoreactivity, supporting a model in which barrier disruption promotes loss of tolerance to brain antigens. At the clinical level, higher baseline autoantibody burden predicts reduced responsiveness to the antipsychotic risperidone, suggesting that autoantibodies contribute to treatment resistance. Together, these findings identify humoral autoimmunity as a pervasive component of schizophrenia and imply that therapies that reset humoral immunity or reduce autoantibody burden may benefit patients beyond those with currently recognized antibody-mediated syndromes.

**GRAPHICAL ABSTRACT:** 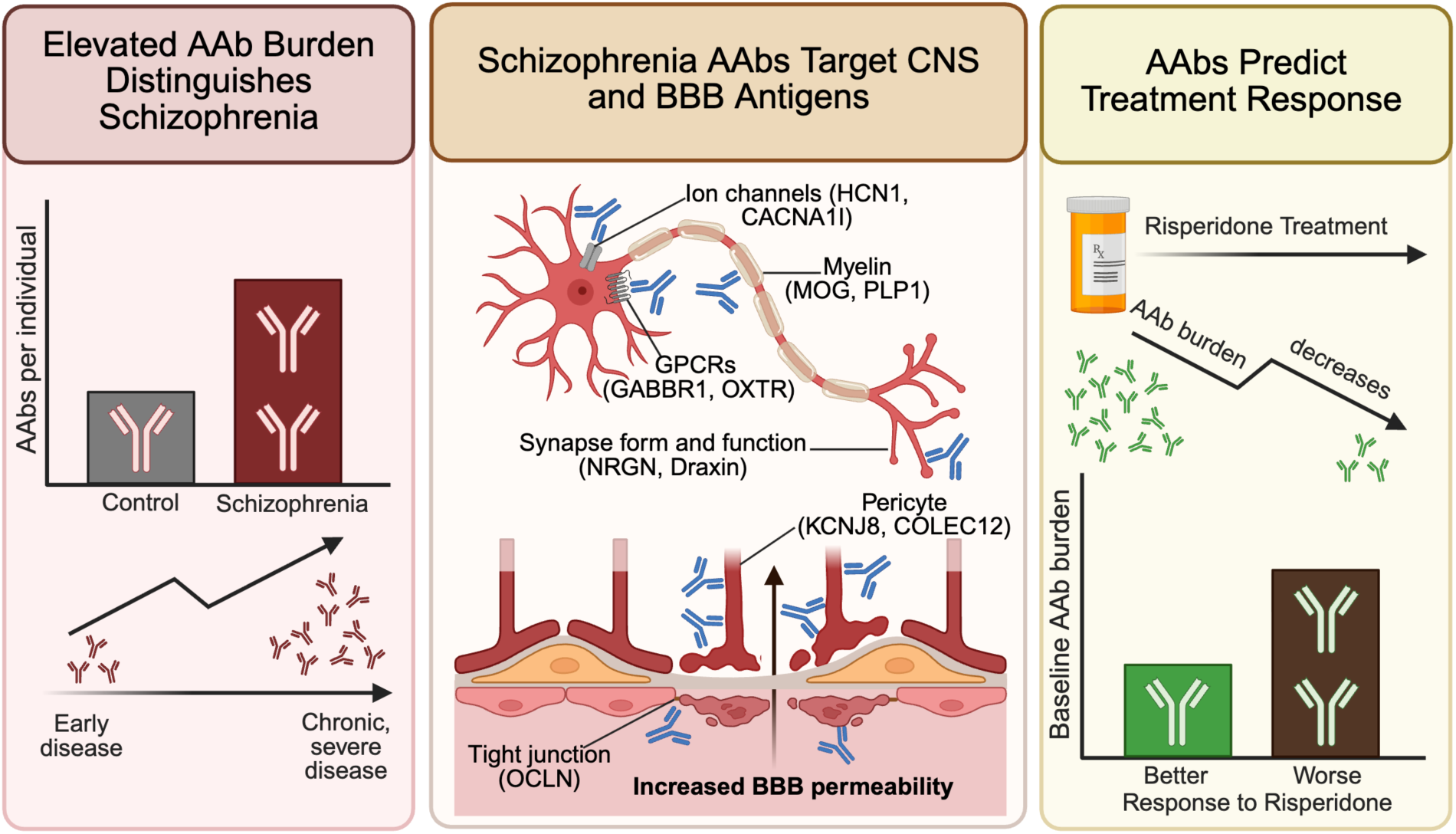

## INTRODUCTION

Schizophrenia is a debilitating neuropsychiatric disorder marked by substantial biological and clinical heterogeneity that remains poorly explained by current models. Current treatments have limited efficacy for negative and cognitive symptoms, and a substantial fraction of patients develop treatment-resistant illness. This heterogeneity suggests multiple underlying biological mechanisms, including immune contributions in a subset of patients. Epidemiological and clinical observations consistent with immune dysregulation include increased susceptibility to severe infection^1,2^, a higher prevalence of autoimmune disease^3^, and altered humoral responses to vaccination^4^. Genetic studies have identified the major histocompatibility complex as the strongest common risk locus in schizophrenia, with additional enrichment of risk at regulatory elements active in CD19+ and CD20+ B cells^5^. Inflammatory abnormalities have also been reported in blood^6^, cerebrospinal fluid^7^ and brain^8^. Yet despite these observations, the role of specific immune-mediated mechanisms in schizophrenia remains incompletely understood. In this context, humoral (antibody-mediated) immunity represents a plausible but underexplored axis of variation^9^. Autoantibodies against extracellular proteins are of particular interest because they can directly modulate neural and neurovascular signaling pathways and can be therapeutically targeted.

Proof of principle for antibody-mediated psychosis comes from autoimmune encephalitides, most notably those associated with autoantibodies against the NMDA receptor (NMDAR)^10^. Affected patients present with prominent psychiatric symptoms that respond dramatically to immunomodulation^11^. However, anti-NMDAR and other defined encephalitis autoantibodies are detected in only a small subset of patients with schizophrenia who lack overt neurological features^12,13^. At the same time, tissue-based assays and cerebrospinal fluid studies suggest that a broader subset of patients harbor antibodies reactive to brain tissue^14,15^, and some exhibit abnormalities consistent with blood–brain barrier dysfunction^16^. Consistent with this possibility, early interventional studies have begun to test immune-directed therapy in psychosis. Immunotherapy has been shown to be feasible and well tolerated in antibody-positive acute psychosis^17^, and B-cell depletion with rituximab has shown potential benefit in treatment-resistant schizophrenia,^18^, motivating randomized trials open to patients regardless of antibody status^19^. Together, these observations raise the possibility that humoral immune mechanisms in schizophrenia extend beyond rare classical autoimmune syndromes and may involve a more extensive repertoire of central nervous system and neurovascular targets. Whether such responses are molecularly structured, clinically meaningful, or relevant to treatment heterogeneity remains unknown.

Here, we hypothesized that schizophrenia is associated with a broader landscape of autoantibodies against extracellular and secreted proteins that contributes to biological and clinical heterogeneity. Such antibody responses could help explain variation in central nervous system autoreactivity, neurovascular barrier integrity, and responsiveness to antipsychotic treatment, while also providing a mechanistic basis for immune-directed therapeutic strategies, including emerging efforts to evaluate B-cell depletion in schizophrenia. To test this hypothesis, we used Rapid Extracellular Antigen Profiling (REAP)^20^ to conduct an autoantibody-wide association study spanning multiple clinically distinct schizophrenia cohorts and a large community-based control cohort without serious mental illness.

## RESULTS

### Elevated autoantibody burden is a core immunologic feature of schizophrenia

To define the landscape of extracellular autoantibodies in schizophrenia, we profiled plasma samples from 352 individuals with schizophrenia and 971 community controls using REAP (**Fig. 1A**). The schizophrenia metacohort comprised distinct sub-cohorts spanning a broad range of disease phenotypes, including antipsychotic-naive (n=62), early-disease (n=43) and chronic psychosis (n=247) patients. The control cohort consisted of community-based individuals without serious mental illness recruited through the NKI Rockland Sample^21,22^. Schizophrenia cases and controls differed in baseline demographics: cases were younger (**Fig. S1A**), more frequently male (**Fig. S1B**), and more frequently reported Black/African American race/ethnicity relative to controls (**Fig. S1D**). Additionally, there were different frequencies of comorbidities between the groups, with schizophrenia cases having a higher incidence of metabolic disease, autoimmune disease, hepatitis, and HIV, consistent with previous reports of these common comorbidities with schizophrenia (**Fig. S1C**)^23,24^.

**Figure 1:**
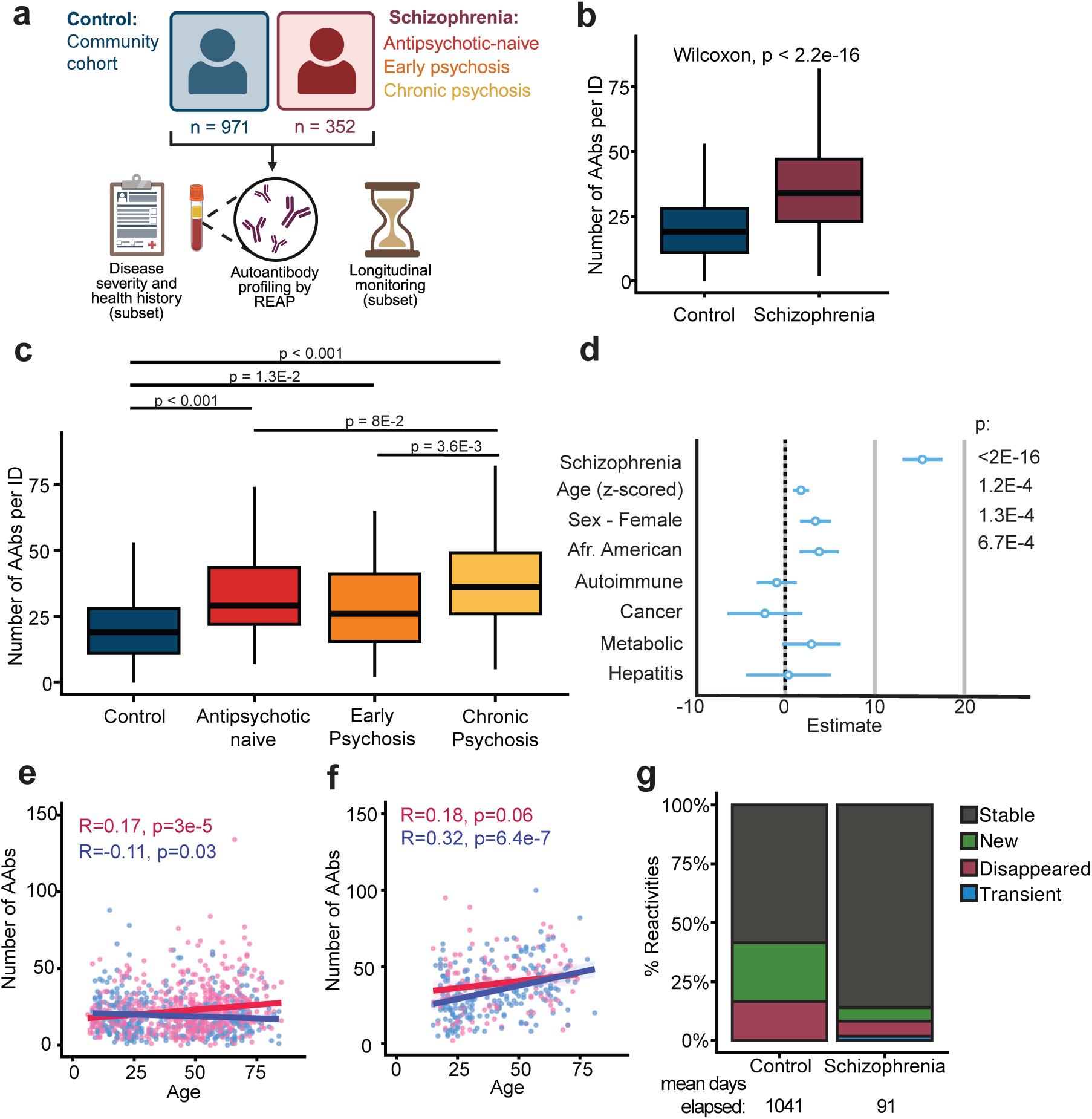
Schizophrenia patients harbor an elevated burden of autoantibodies. **A.** Study overview. **B.** The number of autoantibody (AAb) reactivities per individual (ID) by group (control, n=971; schizophrenia, n = 352). Significance was assessed using unpaired two-sided Wilcoxon test. For the box plots, the central lines indicate the group median values, the top and bottom lines indicate the 75th and 25th percentiles, respectively, the whiskers represent 1.5× the interquartile range. **C.** The number of autoantibody (AAb) reactivities per individual (ID) by group (control, n=971; antipsychotic naive, n = 62; early psychosis, n = 43; chronic psychosis, n = 247). Significance was assessed using Kruskal Wallis (p <2.2E-16) followed by post-hoc Dunn’s multiple comparison tests. P-values were adjusted using the Holm-Bonferroni method. For the box plots, the central lines indicate the group median values, the top and bottom lines indicate the 75th and 25th percentiles, respectively, the whiskers represent 1.5× the interquartile range. **D.** Multivariable linear model predicting the number of autoantibodies per individual as a function of schizophrenia, age (z-scored), sex, race, and comorbid conditions including autoimmune disease, cancer, metabolic disease, and hepatitis. N = 971 control, 264 schizophrenia. Points indicate estimated coefficients and error bars represent 95% confidence intervals. Overall model: p value = <2.2e-16, adjusted R squared: 0.1912. **E, F.** The relationship between number of autoantibody (AAb) reactivities per individual (ID) and age, grouped by sex, among the control population (n = 971) **(E)** and schizophrenia cohort (n=352) **(F)**. Correlation was assessed using Spearman’s correlation. The magenta and blue lines show the linear regression for females and males, respectively, and the shading shows the 95% CIs. Each dot represents one individual (pink = female, blue = male). **G**. Proportion of autoantibody reactivities that changed over time by group (control, subjects = 56, reactivities = 1,009; schizophrenia, subjects = 135, reactivities = 4,692). Stable: autoantibody present at first time point for a given subject (REAP score >=1) is still present at the last time point for that subject (REAP score >= 0.5); New: autoantibody not present at the first time point for a given subject (REAP score <1) is present at the final timepoint for that subject(REAP score >= 1); Disappeared: autoantibody present at first time point for a given subject (REAP score >=1) is not present at the last time point for that subject (REAP score <0.5); Transient: autoantibody not present at first time point for a given subject, appears in intermediate timepoints (REAP score >= 1), and disappears by the last time point (REAP score <0.5).

We found that individuals with schizophrenia harbored a markedly elevated burden of extracellular autoantibodies (**Fig. 1B**). This increase tracked with disease severity and duration, with long-stay psychiatric inpatients exhibiting the highest autoantibody burden—nearly double the number of autoantibody reactivities on average compared to controls (**Fig. 1C**). Individuals with earlier disease, including antipsychotic naive and early psychosis patients, also had elevated autoreactivities at an intermediate level between chronic cases and controls. Overall, we detected reactivity to 3,258 antigens across the metacohort, highlighting the remarkable diversity of the autoantibody reactome in schizophrenia. As we previously observed in other populations^25–28^, these reactivities were distributed across a wide range of frequencies, though most were uncommon and present in <1% of individuals (**Fig. S1E**).

To account for differences in demographics and medical comorbidities between cohorts, we performed multivariable linear regression. Our model confirmed that schizophrenia status was the primary determinant of autoantibody burden **(Fig. 1D)**. Although sex, age, and race showed significant associations, the effect size for schizophrenia (𝛃 = 15.3) was more than four-fold greater than that of any other demographic covariate. We nevertheless examined the relationship between age, sex, and schizophrenia diagnosis on autoantibody levels in greater detail. In control individuals, autoantibody burden increases with age in women but slightly decreases with age in men, with the two trajectories diverging between age 40-80 (**Fig. 1E, S1F**). In contrast, autoantibody burden increases with age in males with schizophrenia (**Fig. 1F, S1G, S1H**), consistent with an estimated gain of approximately 1 additional autoantibody for every 3 years of age. Overall, our findings show that there is a global elevation in autoantibody burden that is present early in the symptomatic period of disease and increases with age in a manner divergent from controls.

To assess autoantibody dynamics in schizophrenia, we tracked individual reactivities in a subset of patients and controls for whom we had longitudinal samples. We observed that autoantibody reactivities in both groups were largely stable over time. Specifically, we found 93% and 78% of individual reactivities observed in schizophrenia and control patients remained present in longitudinal samples spanning an average of approximately 3 months and 3 years, respectively (**Fig. 1G, S1I**). This stability suggests that the age-associated increase in autoantibody burden reflects progressive accrual rather than transient fluctuation.

### Autoantibody diversity and CNS reactivity differentiate schizophrenia from controls

To determine whether individuals with schizophrenia exhibit distinguishable autoantibody signatures relative to controls, we trained a regularized logistic regression (Lasso) classifier (**Fig. 2A**). Under sevenfold cross-validation, the model robustly discriminated schizophrenia cases from community controls without serious mental illness (mean ROC AUC = 0.88). To interpret the features underlying classifier performance, we stratified autoantibody reactivities by their prevalence in the control cohort, defining autoantibodies detected in >1% of controls as “common” and those detected in <1% of controls as “low-frequency.” Schizophrenia cases displayed elevated numbers of both common (**Fig. 2B**) and low-frequency reactivities (**Fig. 2C**).

**Figure 2:**
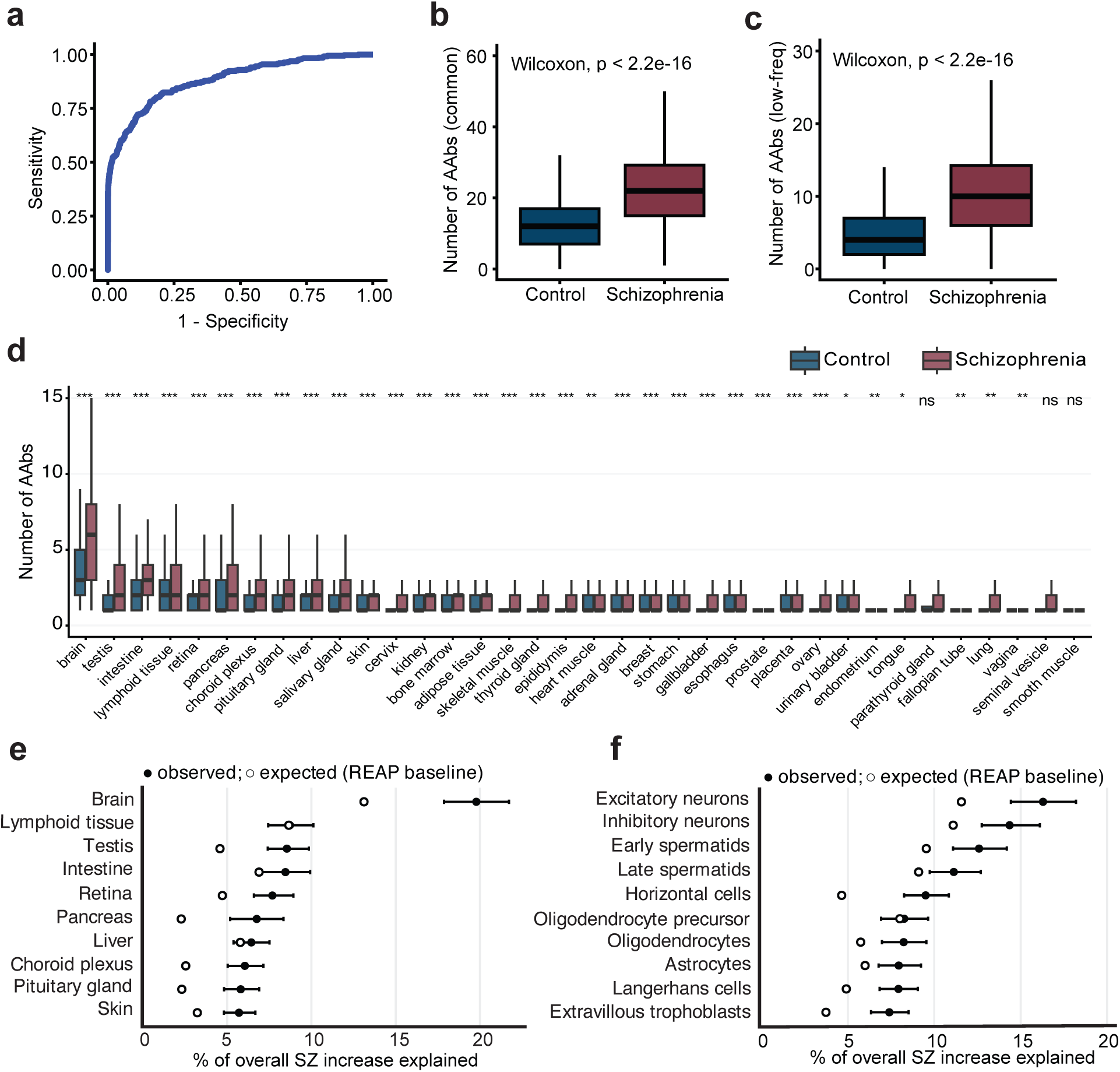
A CNS-directed autoantibody signature distinguishes schizophrenia cases from healthy controls. **A.** Receiver Operator Classification (ROC) curve of the REAP-Lasso model for patient vs. control discrimination (AUC = 0.884). **B.** The number of common autoantibody (AAb) reactivities per individual by group (control, n=971; schizophrenia, n = 352). Common reactivities defined as present in >1% of the control cohort. Significance was assessed using unpaired two-sided Wilcoxon test. For the box plots, the central lines indicate the group median values, the top and bottom lines indicate the 75th and 25th percentiles, respectively, the whiskers represent 1.5× the interquartile range. **C.** The number of low-frequency autoantibody (AAb) reactivities per individual by group (control, n=971; schizophrenia, n = 352). Low-frequency reactivities defined as present in <= 1% of the control cohort. Significance was assessed using unpaired two-sided Wilcoxon test. For the box plots, the central lines indicate the group median values, the top and bottom lines indicate the 75th and 25th percentiles, respectively, the whiskers represent 1.5× the interquartile range. Low-freq = Low-frequency **D.** The number of autoantibody (AAb) reactivities against each tissue category per individual by group (control, n = 971; schizophrenia = 352). Tissue categories are composed of REAP reactivities bucketed by human protein atlas mRNA expression data. Significance was assessed by unpaired two-sided Wilcoxon with correction for multiple hypotheses by Benjamini-Hochberg. *** ∼ <0.001, ** ∼ <0.01, * ∼ <0.05. **E, F.** Forest plots depicting the top tissues **(E)** or single cell types **(F)** (y-axis) ranked by their estimated contribution to the overall schizophrenia vs. control increase in total autoantibodies (AAbs), with the x-axis reporting percentage of overall schizophrenia increase explained. Tissue or single cell categories are composed of REAP reactivities bucketed by human protein atlas mRNA expression data. Because a given protein may belong to more than one tissue class, percentages need not sum to 100. Closed circles (●) represent the observed percent contribution. Open circles (○) indicate the expected contribution under a REAP-library baseline. Horizontal whiskers denote 95% confidence intervals from a cohort-/group-stratified bootstrap that resamples subjects within schizophrenia and control groups (R = 4000 replicates).

To assess whether specific tissues or cell types are preferentially targeted in schizophrenia, we annotated REAP antigens by tissue and cell-type expression using Human Protein Atlas (HPA) mRNA data. We then quantified both the raw number of reactivities and the fraction of the overall increase in autoantibodies in schizophrenia attributable to each tissue or cell-type category. Autoantibodies against brain-associated antigens accounted for the largest single tissue category, with schizophrenia patients possessing more than six brain-directed autoreactivities on average (**Fig. 2D**). Brain autoantibodies also represented roughly one-fifth of the overall excess autoantibody signal in schizophrenia patients relative to controls (**Fig. 2E**). At the cell-type level, excitatory neurons and inhibitory neurons each explained a substantial share of both the raw autoantibody burden (**Fig. S2A**) and the excess antibody signal (16% and 14%, respectively) (**Fig. 2F**), indicating that the schizophrenia-associated autoantibody increase is disproportionately neuronal. Testis-related cell types (early/late spermatids) and other brain cell types (*e.g.*, oligodendrocytes and astrocytes) also contributed meaningfully to a lesser extent. Taken together, these results define a schizophrenia-associated autoantibody signature consisting of both common and rare autoantibody reactivities that are weighted towards central nervous system antigens, suggesting a disease-specific pattern of humoral immune dysregulation.

### Schizophrenia-associated autoantibodies bind diverse brain- and neuron-enriched proteins

Beyond global autoantibody signatures, we next asked whether specific autoreactivities were enriched in schizophrenia. We therefore conducted an autoantibody-wide association study (AAbWAS), analogous to a GWAS, in which we calculated odds ratios for each detected autoreactivity by comparing its occurrence in schizophrenia cases versus controls. Because age, sex and race influenced autoantibody burden, odds ratios were adjusted for these variables. This analysis identified 93 autoantibodies enriched in schizophrenia with an odds ratio greater than 2.72 (estimate >1) and a q-value less than 0.05 (**Table S1**). On average, individuals with schizophrenia possessed approximately five schizophrenia-enriched reactivities, with some harboring up to 20 such autoantibodies (**Fig. S3A**).

Because brain and neuronal compartments accounted for a disproportionate share of the overall autoantibody increase (**Fig. 2D,E,F, S2A**), we next focused on schizophrenia-enriched autoantigens with elevated expression in the brain, excitatory neurons, or inhibitory neurons according to HPA mRNA data. This analysis highlighted 26 schizophrenia-enriched brain- and neuron-associated autoantigens (**Fig. 3A,B**), spanning diverse functional classes that included ion channels involved in neuronal excitability (**Fig. 3C,D**), proteins linked to synaptic function (**Fig. 3E**), extracellular matrix and myelin components (**Fig. S3B**), and neuromodulatory G-protein-coupled receptors (GPCRs) (**Fig. 3F**).

**Figure 3:**
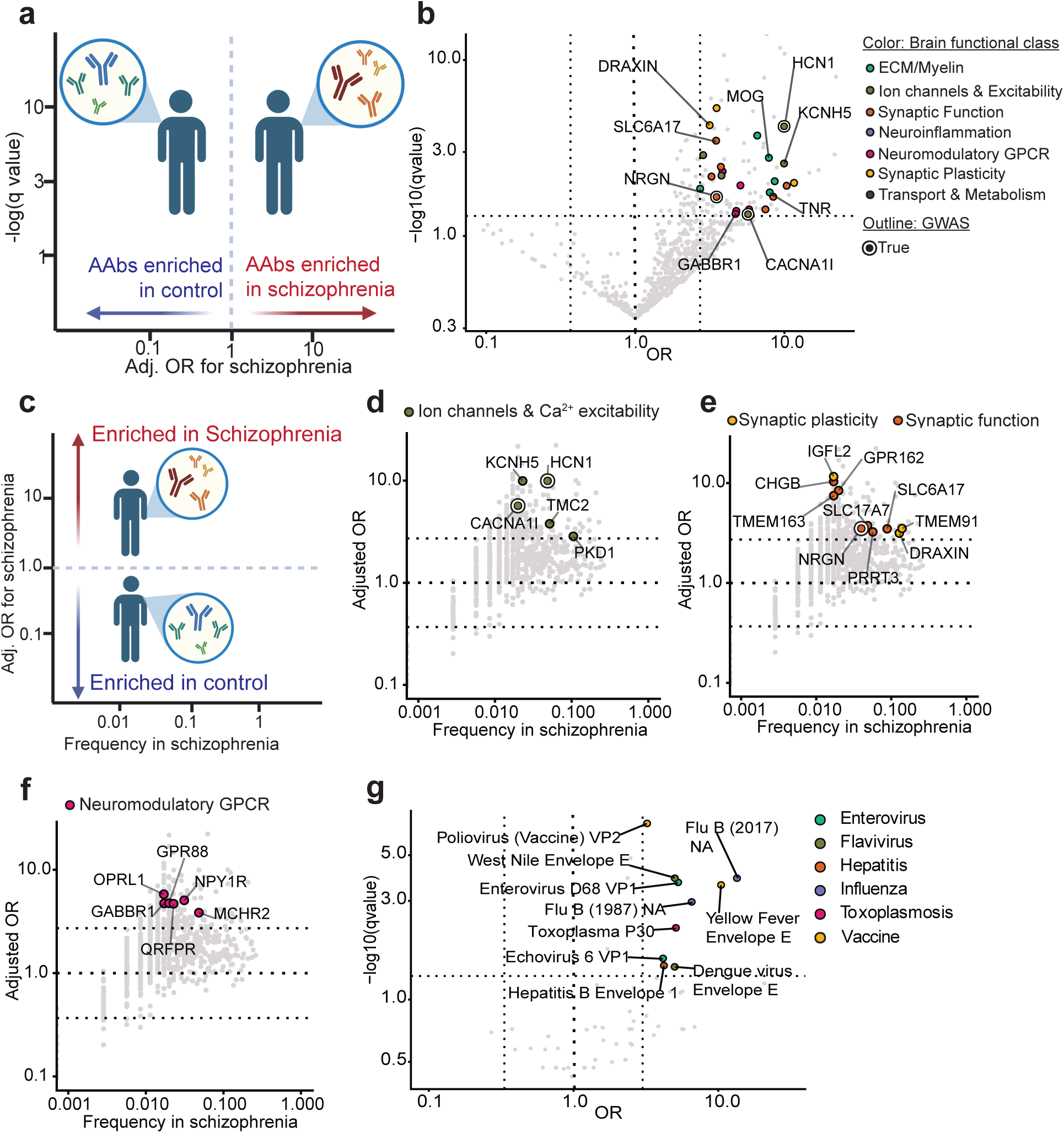
Schizophrenia-enriched autoantibodies target proteins critical for neuronal excitability and synaptic function. **A.** Conceptual figure for depiction of autoantibody odds ratio for schizophrenia status. **B.** Volcano plot depicting all autoantibody reactivities detected. Reactivities that are significantly enriched in schizophrenia (q-value <0.05, odds ratio (OR) >2.72) and target proteins with elevated expression in the brain and inhibitory and excitatory neurons by human protein atlas mRNA expression are bolded and colored. Color indicates functional annotation. Each dot represents one autoantibody reactivity. Vertical dashed lines indicate OR threshold +/-2.72. Horizontal dashed line indicates q-value threshold of 0.05. **C.** Conceptual figure for depiction of autoantibody odds ratio for schizophrenia status and AAb frequency in schizophrenia. **D,E,F.** Frequency plot depicting all autoantibody reactivities detected, with proteins belonging to the ion channel and Ca+2 excitability group **(D)**, synaptic function and synaptic plasticity groups **(E)**, and neuromodulatory GPCR **(F)** from figure B bolded and colored. Horizontal dashed lines indicate OR threshold of +/-2.72. Each dot represents one reactivity. **G.** Volcano plot depicting all anti-pathogen reactivities detected. Reactivities that are significantly enriched in schizophrenia (q-value <0.05, odds ratio (OR) >2.72) are bolded and colored. Color indicates pathogen class annotation. Each dot represents one reactivity. Vertical dashed lines indicate OR threshold +/-2.72. Horizontal dashed line indicates q-value threshold of 0.05

Five neuronal ion channels were disproportionately targeted in schizophrenia (**Fig. 3D**), and these reactivities were collectively common, with approximately one quarter of patients harboring at least one ion-channel autoantibody. Particularly notable were reactivities against HCN1, a cation channel that regulates cortical resting membrane potential, whose binding was validated by ELISA (**Fig. S3C**), and CACNA1I (Cav3.3), a T-type calcium channel essential for thalamic function; both have been identified as schizophrenia risk loci by GWAS^5^. Electrophysiological recordings in CACNA1I-expressing HEK293 cells showed numerically lower current density in cells treated with IgG from CACNA1I-autoantibody-positive patients, though the difference did not reach statistical significance (**Fig. S4A-E**). Additional schizophrenia-enriched brain autoantigens included proteins involved in synaptic and circuit function, such as NRGN, which regulates calcium–calmodulin-dependent synaptic plasticity and has also been implicated by schizophrenia GWAS^29^, and Draxin, an axon-guidance factor important for neural circuit formation^30^; approximately 10% of schizophrenia patients possessed Draxin autoantibodies (**Fig. 3E**).

Schizophrenia-enriched autoreactivities also targeted structural components that support neuronal function, including the extracellular matrix protein TNR and the myelin proteins MOG and PLP1, with nearly 10% of patients harboring PLP1 autoantibodies (**Fig. S3B**). We identified autoantibody reactivities against multiple neuromodulatory GPCRs (**Fig. 3F).** These included GPR88, a striatal receptor that modulates dopamine D2 receptor signalling, and GABBR1, a subunit of the GABA-B receptor that helps set inhibitory tone in the brain. GABBR1 is of particular clinical interest because GABA-B receptor autoantibodies have been previously detected in cases of autoimmune encephalitis associated with psychotic symptoms^31–33^.

Additional GPCR targets included NPY1R, QRFPR, and MCHR2, which have established roles in hypothalamic control of arousal, stress, feeding, and affect. Consistent with the prominence of neuroactive receptors among the schizophrenia-enriched autoantigens, KEGG pathway analysis identified the Neuroactive Ligand–Receptor Interaction pathway as the most enriched, though this did not survive multiple testing correction (hsa04080; fold enrichment = 2.1, q = 0.12; **Fig. S3D**). Several of these proteins have been implicated in neuroinflammation and neurovascular signaling, including HRH4, C3AR1, BDKRB2, and TBXA2R **(Fig. S3E**). HRH4, a histamine receptor that influences microglial migration and cytokine release, was one of the most differentially targeted proteins, with autoreactivity detected in 22 of 352 schizophrenia cases compared with 3 of 972 controls. We also detected autoreactivity against OXTR, which regulates social cognition and stress responsivity, though OXTR autoantibodies did not inhibit receptor signaling in an *ex vivo* β-arrestin recruitment assay (**Fig. S3F,G**).

### Schizophrenia-associated antibody signatures extend to pathogen antigens

Because the REAP library also includes selected pathogen antigens, we next asked whether pathogen-directed antibody responses differed between schizophrenia and controls **(Fig. 3G**). Schizophrenia patients displayed increased reactivity to *Toxoplasma gondii* P30/SAG1, the parasite’s major tachyzoite surface antigen, consistent with prior work linking *T. gondii* seropositivity to schizophrenia^34^. We also observed enriched reactivity to a cluster of neurotropic flavivirus envelope antigens, including yellow fever, West Nile, and dengue viruses, as well as to poliovirus VP2, enterovirus D68, echovirus 6 VP1, influenza B neuraminidase, and hepatitis B envelope 1. Although schizophrenia-specific associations for several of these pathogens remain uncertain, the convergence on neuroinvasive viruses is notable given epidemiologic evidence linking CNS viral infection to later psychotic illness^35,36^.

### Schizophrenia-associated autoantibodies target neurovascular barrier components and impair BBB integrity

Given prior reports implicating altered blood–brain barrier (BBB) and blood–cerebrospinal fluid (B-CSF) barrier function in schizophrenia^37^, we asked whether schizophrenia-enriched autoantibodies targeted proteins present in these structures. The BBB is formed by brain endothelial cells through interactions with pericytes, astrocytes, and vascular smooth muscle cells, whereas the B-CSF barrier is primarily formed by choroid plexus epithelial cells. Using Human Protein Atlas expression data, we found that 14 of 93 schizophrenia-enriched autoantibodies recognized proteins with elevated expression in one or more of these five cell types (**Fig. 4A**). These included autoantibodies against OCLN, a critical tight junction protein present at the BBB, which was targeted in 15 individuals with schizophrenia. We also identified reactivities against proteins associated with pericytes and vascular smooth muscle cells, including COLEC12, a lipid scavenger receptor, and KCNJ8, an ATP-sensitive potassium channel important for both pericyte and vascular smooth muscle development and neurovascular coupling. Additional schizophrenia-associated reactivities targeted barrier-associated transporters, including SLC52A3, a riboflavin (vitamin B2) transporter and SLC2A2, a low-affinity glucose transporter.

**Figure 4:**
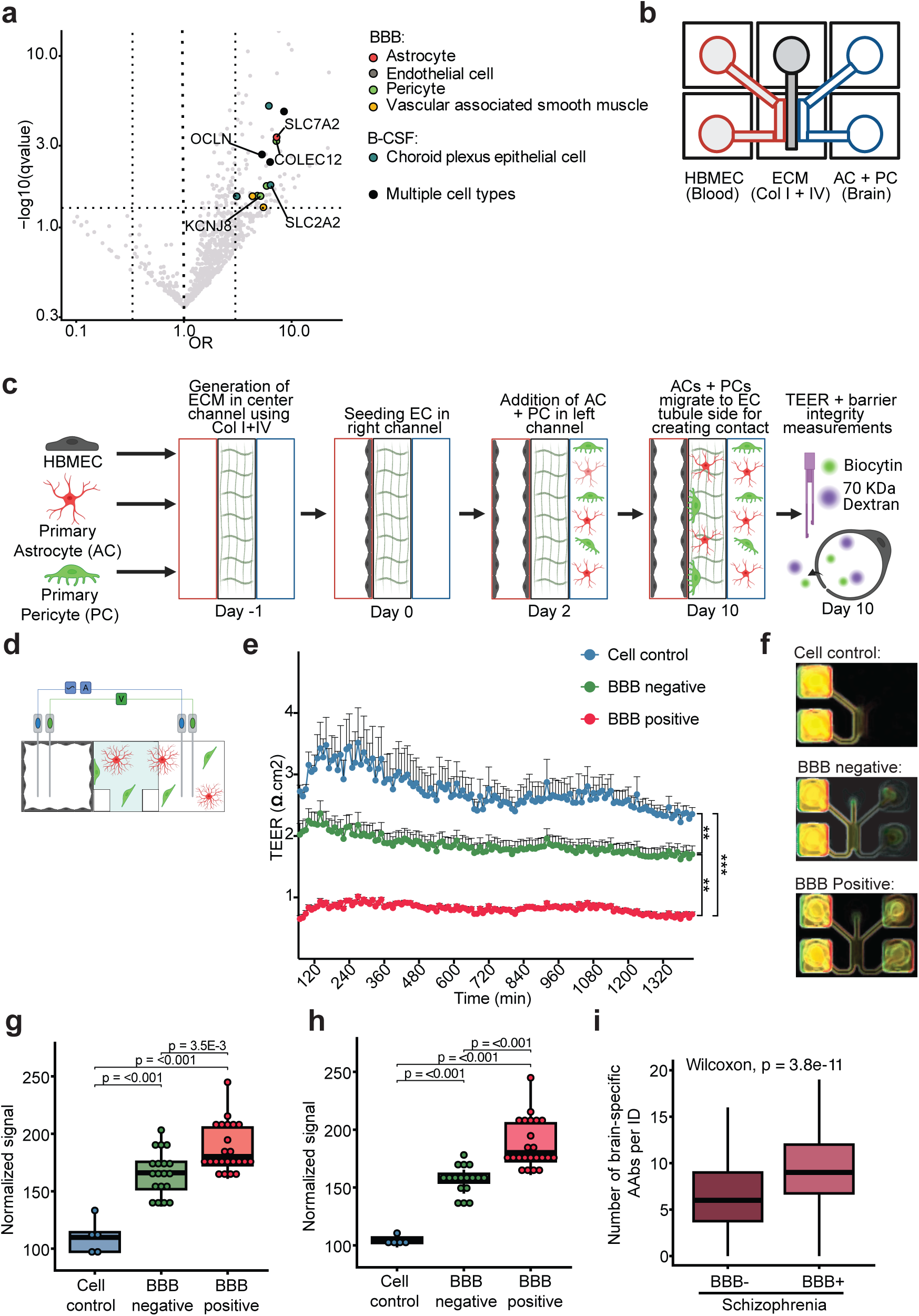
Functional autoantibodies target the blood-brain and blood-CSF barriers in schizophrenia. **A.** Volcano plot depicting all autoantibody reactivities detected, with reactivities that are significantly enriched in schizophrenia (q-value <0.05, odds ratio (OR) >2.71) and target proteins elevated in brain pericytes, endothelial cells, astrocytes, vascular associated smooth muscle, or choroid plexus epithelial cells (by HPA mRNA expression data) bolded and colored. Each dot represents one autoantibody reactivity. Vertical dashed lines indicate OR threshold +/-2.71. Horizontal dashed line indicates q-value threshold of 0.05. **B.** Graphic representation of the MIMETAS chip used for the 3D human BBB model with flow. **C.** Diagram illustrating the creation of the 3D human BBB model and experimental scheme for the tracer permeability assay. **D.** Graphic representation of the experimental scheme for the TEER measurement with the OrganoTEER instrument. **E.** Graph of the TEER measurements over time in the 3D human BBB model under three conditions. Each dot represents one measurement. ** p<0.01, **** p<0.001. **F.** Representative images of tracer leakage from the blood (left) into the brain (right) compartment of the 3D BBB model. Biocytin is in green and Dextran is in red. There is increased tracer in the brain compartment with the BBB positive sera from Schizophrenia patients. **G,H.** Plots of the normalized signal of biocytin **(G)** and 70 KDa Dextran **(H)** permeability measurements at 30 minutes after tracer application. Significance was assessed using one way ANOVA (p=3.2E-9, 2.6E-12) followed by Tukey’s post hoc test. For the box plots, the central lines indicate the group median values, the top and bottom lines indicate the 75th and 25th percentiles, respectively, the whiskers represent 1.5× the interquartile range. BBB = Blood brain barrier. **I.** Number of autoantibodies (AAb) per individual (ID) targeting proteins with elevated expression in the brain by human protein atlas mRNA expression data. N=328 for BBB-, 24 for BBB+. Significance was assessed using unpaired Wilcoxon. For the box plots, the central lines indicate the group median values, the top and bottom lines indicate the 75th and 25th percentiles, respectively, the whiskers represent 1.5× the interquartile range. BBB = Blood-brain barrier.

To test whether schizophrenia-associated autoreactivity against BBB-related proteins altered endothelial barrier function, we evaluated plasma from schizophrenia patients positive for selected schizophrenia-enriched autoantibodies against BBB-expressed or BBB-regulatory proteins (BBB+) in a 3D microfluidic BBB model consisting of brain endothelial cells, extracellular matrix (ECM), pericytes, and astrocytes layered in a multi-chambered chip^38,39^(**Fig. 4B,C**). As a comparator, we analyzed plasma from schizophrenia patients lacking these reactivities (BBB−). BBB+ plasma caused a greater reduction in trans endothelial electrical resistance (TEER), a readout of paracellular BBB function, than either no-plasma controls or BBB− plasma, indicating impaired BBB paracellular integrity (**Fig. 4D,E**). Consistent with this result, orthogonal tracer assays showed that BBB+ plasma increased permeability to both a small (biocytin, 890 Da) and large (70-kDa dextran) tracer relative to BBB− plasma, indicating increased paracellular and transcellular permeability (**Fig. 4F–H**).

Given the central role of the BBB in maintaining immune privilege and limiting exposure of brain antigens to the peripheral immune system^40^, we next asked whether schizophrenia patients harboring BBB-associated autoantibodies exhibited broader central nervous system autoreactivity. Individuals with at least one BBB autoantibody reactivity displayed nearly twice as many autoantibody reactivities against HPA brain-elevated proteins as schizophrenia cases lacking these reactivities (**Fig. 4I**).

### Autoantibody burden is associated with antipsychotic responsiveness and declines during risperidone treatment

We next asked whether schizophrenia-associated autoantibodies were associated with response to antipsychotic treatment. We therefore profiled the autoantibody reactomes of a cohort of antipsychotic-naive patients with schizophrenia during a treatment trial of risperidone (**Fig. 5A**). As expected, symptoms improved over the course of the 16-week study, as assessed by the Positive and Negative Symptom Scale for Schizophrenia (PANSS) (**Fig. 5B**). Improvements were observed in both positive and general psychopathology subscales (**Fig. 5C, S5A,B**), whereas negative symptom subscales did not change significantly at the cohort level (**Fig. 5C, S5C**).

**Figure 5:**
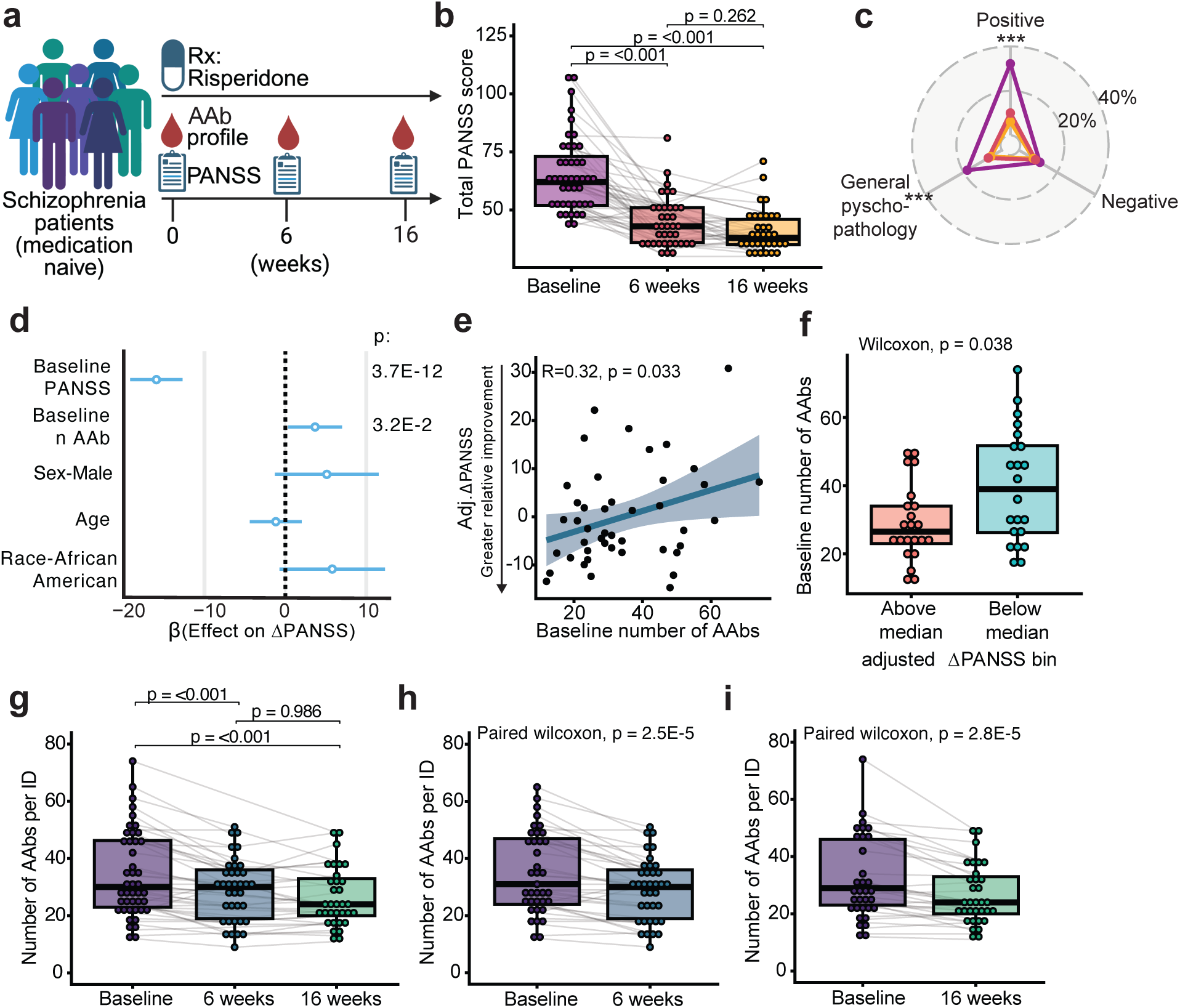
Autoantibody burden is modified by the antipsychotic risperidone and predicts treatment response. **A**. Study overview. **B.** Total PANSS score for participants (n = 44) during the study. Data were modeled using a linear mixed-effects model (n=44 patients) with timepoint as a fixed effect and subject identity as a random intercept. The overall effect of time was assessed via a Type III Wald F-test with Kenward-Roger degrees of freedom approximation (F(2, 74.0) = 54.1, p = 3.3E-15). Pairwise comparisons were performed using estimated marginal means with p-values adjusted for multiple comparisons using the Holm-Bonferroni method. Each dot represents one individual per time point, with multiple time points for a given individual connected by grey lines. For the box plots, the central lines indicate the group median values, the top and bottom lines indicate the 75th and 25th percentiles, respectively, the whiskers represent 1.5× the interquartile range. **C.** Mean PANSS subscores for all study participants (n=44) per timepoint. Purple = baseline; orange = 6 weeks; yellow = 16 weeks. Percentage scale indicates percentage of max score. Data were modeled using a linear mixed-effects model (n=44 patients) with timepoint as a fixed effect and subject identity as a random intercept. The overall effect of time was assessed via a Type III Wald F-test with Kenward-Roger degrees of freedom approximation (Positive: F(2, 74) = 955.5, p = <2.2E-16), (Psychopathology: F(2, 74.9) = 45.0, p = 1.4E-13), (Negative: F(2, 72.3) = 1.4, p = 0.25). **D.** Multivariable linear model predicting the total PANSS delta during the study as a function of baseline PANSS score, baseline number of autoantibody (AAb) reactivities, age (z-scored), sex, and race. N = 44. Points indicate estimated coefficients and error bars represent 95% confidence intervals. Overall model: p value = 8.84e-10, adjusted R squared: 0.6903. **E.** Relationship between baseline autoantibody (AAb) burden and change in symptom severity after adjustment for covariates. The y-axis reflects residualized values from a linear model in which change in PANSS score (ΔPANSS) was adjusted for baseline PANSS score, age, race, and sex. Points represent individual subjects (n = 44), and the line indicates the fitted linear association (spearman) between adjusted ΔPANSS and baseline number of autoantibodies with 95% CI shaded. **F.** baseline number of autoantibodies (AAbs) per individual, with individuals bucketed by change in PANSS score (ΔPANSS) after adjustment for covariates (baseline PANSS, age, race, and sex). Statistical significance assessed by unpaired two-sided Wilcoxon. Each point represents one individual (n=44). **G.** Number of autoantibody (AAb) reactivities per individual (ID) at baseline, 6 weeks, and 16 weeks of risperidone treatment. Data were modeled using a linear mixed-effects model (n=44 patients) with timepoint as a fixed effect and subject identity as a random intercept. The overall effect of time was assessed via a Type III Wald F-test with Kenward-Roger degrees of freedom approximation (F(2, 68.8) = 22.0, p = 4.2E-8). Pairwise comparisons were performed using estimated marginal means with p-values adjusted for multiple comparisons using the Holm-Bonferroni method. For the box plots, the central lines indicate the group median values, the top and bottom lines indicate the 75th and 25th percentiles, respectively, the whiskers represent 1.5× the interquartile range. **H, I.** Number of autoantibodies (AAbs) per individual (ID) for patients with paired timepoints for 0 to 6 (**H**) and 0 to 16 (**I**) (n = 38, n = 39, respectively). Statistical significance was determined by paired two-sided Wilcoxon. For the box plots, the central lines indicate the group median values, the top and bottom lines indicate the 75th and 25th percentiles, respectively, the whiskers represent 1.5× the interquartile range.

To evaluate the relationship between baseline autoantibody burden and therapeutic response (ΔPANSS), we developed a multivariable linear regression model. After adjusting for baseline PANSS, age, sex, and race, our analysis revealed that the number of baseline autoantibody reactivities significantly predicted treatment outcomes, with a higher autoantibody burden associated with decreased symptom reduction (**Fig. 5D,E**). To further visualize this association, we stratified patients by relative symptom improvement (ΔPANSS adjusted for baseline PANSS). Patients with above-median improvement had significantly fewer autoantibody reactivities at baseline than those with below-median improvement (**Fig. 5F**).

Given the association of baseline autoantibody burden with treatment response, we next asked whether the autoantibody reactome changed over the course of risperidone treatment. Longitudinal analysis showed that the number of autoantibody reactivities per individual declined significantly during treatment (**Fig. 5G, H, I**). This decrease extended to autoantibodies against both the previously defined sets of brain and neuron-enriched proteins and schizophrenia-enriched antigens (**Fig. S5D,E**). Together, these findings indicate that elevated baseline autoantibody burden is associated with reduced antipsychotic responsiveness and that risperidone treatment is associated with a decline in autoantibody burden.

## DISCUSSION

Immune mechanisms have long been implicated in schizophrenia, yet the relevance of humoral immunity has remained uncertain outside rare autoimmune encephalitides. Here, we identify a striking elevation in extracellular autoantibody burden in schizophrenia, with preferential targeting of central nervous system and neurovascular antigens. These responses were detectable in early-stage cohorts, were most pronounced in chronic psychosis and, for many individual autoantibodies, remained stable over time. Together, these findings establish autoantibody reactivity as a broad rather than exceptional feature of schizophrenia, spanning diverse disease-relevant targets.

A notable feature of the schizophrenia autoantibody reactome is its convergence on molecules with established genetic and mechanistic links to schizophrenia, including HCN1, CACNA1I and NRGN^5,29^. This overlap suggests that humoral and genetic variation may intersect at shared pathways governing neuronal excitability, synaptic function and circuit regulation. At the same time, not all schizophrenia-associated autoantibodies produced detectable functional effects in our assays, including those targeting CACNA1I and OXTR. Some may reflect impaired B cell self-tolerance, functioning as biomarkers rather than direct effectors of disease. Others may act through mechanisms not captured by the target-focused assays we applied, including Fc receptor- and complement-mediated effector functions. Consistent with this possibility, recent preclinical work in an anti-NMDAR autoimmunity model showed that FcRγ-chain-deficient mice failed to develop the full psychosis-like phenotype despite generating comparable anti-NMDAR autoantibody titers, implicating Fc receptor engagement as a determinant of organism-level pathology^41^. Importantly, the absence of robust effects in simplified systems does not exclude biological relevance: even modest perturbations in receptor or channel function may be sufficient to alter neural circuit dynamics. For example, reductions in CACNA1I current density of ∼20% are sufficient to disrupt thalamic oscillatory activity^42^. More broadly, these findings are consistent with models of schizophrenia in which modest perturbations across convergent neural pathways can give rise to network dysfunction, analogous to the disorder’s highly polygenic architecture. The schizophrenia-associated autoantibody reactome therefore appears diverse rather than restricted to a small set of canonical reactivities, consistent with multiple distinct humoral responses contributing to disease biology.

BBB dysfunction has been implicated in schizophrenia, and the neurovascular targets identified here provide a potential mechanistic link^37^. Plasma from patients with autoreactivity to diverse BBB-associated antigens increased barrier permeability in a human 3D microfluidic BBB model, providing functional evidence that schizophrenia-associated autoantibodies can compromise neurovascular integrity. Some antibodies may perturb the function of their cognate barrier proteins directly, whereas others may converge through shared Fc-dependent inflammatory signalling once localized to the neurovascular surface. Brain endothelial and perivascular cells express Fc receptors, raising the possibility that antibody docking at distinct antigens expressed by BBB-forming cells triggers a more stereotyped downstream program of BBB breakdown. The observation that patients with BBB-directed autoreactivity exhibited broader CNS-directed autoreactivity is consistent with a feed-forward model in which endothelial barrier dysfunction increases peripheral exposure to antigens within an otherwise immune-privileged site, thereby broadening CNS-directed autoreactivity. We also identified antibodies targeting B-CSF barrier. Although not tested here, it is also possible that B-CSF dysfunction may contribute to antibody entry into the cerebrospinal fluid where it would also gain access to several brain antigens.

At the clinical level, autoantibody burden was associated with antipsychotic responsiveness. Higher baseline autoantibody burden predicted reduced responsiveness to risperidone, and autoantibody burden declined during treatment. These observations raise the possibility that humoral immune processes track clinically meaningful variation in antipsychotic response and that risperidone treatment may influence autoantibody burden, directly or indirectly. Autoantibody burden may itself contribute to treatment resistance, or it may instead mark other disease processes, such as broader neuroinflammation or neurovascular dysfunction, that are less responsive to risperidone. The mechanism underlying the treatment-associated decline in autoantibody burden remains unclear. One possibility is that it reflects attenuation of inflammatory signals that sustain autoreactive B cell responses, in line with reports that antipsychotic treatment can reduce inflammatory cytokine levels^43^ and normalize elevated B cell frequencies in acute psychosis.^44^ More direct effects are also plausible, as B cells express dopamine receptors targeted by antipsychotic drugs^45^. Consistent with this possibility, antipsychotic therapies including risperidone and clozapine have been associated with reductions in serum immunoglobulin levels^46,47^, and recent preclinical work further showed that clozapine can reduce disease-relevant anti-NMDAR autoantibody responses in a mouse model of antibody-mediated psychosis^41^. Although these observations do not establish causality, they are consistent with a broader role for disease-relevant autoantibodies in shaping therapeutic response, as we recently observed in cancer immunotherapy^28^.

This study has important limitations. We profiled plasma rather than cerebrospinal fluid and therefore cannot determine which autoreactivities access the CNS or are produced intrathecally. Most identified autoantibodies were not directly functionally tested, and the *ex vivo* BBB findings do not establish the same mechanism *in vivo*. In addition, although the autoantibody signal was evident in earlier-disease cohorts, a substantial fraction of the schizophrenia cohort consisted of institutionalized patients with chronic psychosis, so contributions from environmental exposures, comorbidities and treatment history cannot be fully excluded. Larger independent cohorts will be needed to refine disease-associated autoreactivities, compare the schizophrenia autoantibody reactome relative to other psychiatric conditions, and define potential immune-stratified subgroups. Interventional trials will be required to determine whether therapies that reduce autoantibody burden, deplete antibody-producing cells or more broadly reset humoral immunity improve outcomes.

## METHODS

### Study participants and ethics approval

Plasma samples were obtained from four independent cohorts. All studies were conducted in accordance with the Declaration of Helsinki. All samples were stored at −80°C until analysis. Clinical data elements were de-identified, and all protected health information was removed in accordance with the Health Insurance Portability and Accountability Act.

#### Healthy control cohort

Samples were obtained from the Nathan Kline Institute-Rockland Sample (NKI-RS), a community-ascertained lifespan cohort from Rockland County, New York.^21,22^ Participants were recruited through zip code-based community outreach including targeted mailings, community events, and word-of-mouth referral. Participants with any history of psychotic disorder, bipolar disorder, or other serious mental illness were excluded. The NKI-RS was approved by the Institutional Review Board of the Nathan Kline Institute. All participants provided written informed consent, including consent for data sharing. Access to NKI-RS samples and data was governed by a Data Usage Agreement with the New York State Office of Mental Health.

#### Antipsychotic-naïve first episode psychosis (FEP) cohort

Samples were collected from a cohort of medication-naïve FEP subjects who participated in a 16-week trial of risperidone (NCT03442101). Subjects were recruited from the emergency room inpatient units, and outpatient clinics at the University of Alabama (UAB) at Birmingham. PANSS^48^ score was administered at baseline, week 6, and week 16 post-risperidone treatment. Approval for this study was obtained from the UAB Institutional Review Board and subjects provided written informed consent for use of their data in future studies.

#### Early psychosis cohort

Samples were obtained from a cohort of subjects with early-phase schizophrenia stabilized on antipsychotic medication (<24 weeks of treatment) who participated in a randomized placebo-controlled trial of citalopram (NCT01041274). Participants were recruited from early psychosis clinical programs at New York University Medical Center (New York, NY) and Massachusetts General Hospital/Harvard Medical School (Boston, MA). Approval was obtained from the Institutional Review Board at each site. All participants provided written informed consent, including use of data in future studies.

#### Chronic psychosis cohort

Samples were collected from adult inpatients with schizophrenia and schizoaffective disorder hospitalized in psychiatric centers operated by the New York State Office of Mental Health. Samples were derived from blood collected for routine clinical purposes, and covariate data (including primary psychiatric diagnosis, age, sex, self-reported race/ethnicity, smoking status, and medical conditions) were extracted from electronic health records. The study was approved by the Nathan Kline Institute Institutional Review Board with a waiver of informed consent for secondary use of biospecimens.

### Medical comorbidity classification

Patients were assigned a binary value (yes or no) for all medical condition groups (autoimmune, cancer, hepatitis, HIV, metabolic disease, cardiovascular disease) based on medical records and listed diagnosis terms. Because each cohort originated from a different parent study, diagnosis terms in the clinical data differed. Diagnosis terms from each study that were used to classify a patient as having one of the six medical conditions used in this study are listed in **Supplementary Table 2.**

### Rapid extracellular antigen profiling

#### Antibody purification and yeast depletion

IgG purification was performed as previously described. 20 μL of PBS-washed protein G magnetic resin (Lytic Solutions) were then mixed with 25 μL inactivated plasma and the mixture was incubated at 4 °C for three hours with agitation. The resin was then washed with sterile PBS and resuspended in 90 μL of 100 mM glycine (pH 2.7). Following a 5-minute incubation at room temperature, the supernatant was separated and mixed with 10 μL of 1M Tris (pH 8.0). IgG concentration was then determined using a NanoDrop 8000 Spectrophotometer (Thermo Fisher Scientific). Purified IgG was then combined with 10^8^ induced yeast cells (with empty pDD003 vector) in 100 μL PBE (PBS with 0.5% BSA and 0.5 mM EDTA). After a 3-hour incubation at 4 °C with shaking, the mix was filtered through 96-well 0.45-μm plates (Thomas Scientific) with 3,000g for 3 minutes to collect the yeast-depleted IgG.

#### Yeast library-Antibody selection

The human exoproteome yeast library^28^ and REAP screening method was previously described^20,25^. Briefly, the yeast library was grown in SDO-Ura at 30 °C to achieve OD between 5-7. The library was then induced in 1:10 SDO-Ura:SGO-Ura at 30 °C with starting OD at 1 for 20 hours. Before selection, plasmid DNA was isolated from 400 µL of the induced library (Zymoprep Yeast Cell Plasmid Miniprep II kit) as the baseline reference for antigen frequency. 10 μg yeast-depleted IgG was incubated with 10^8^ induced library yeast in 100 μL PBE with shaking for 1 hour at 4 °C, followed by 30 minutes incubation with 1:100 biotin anti-human IgG Fc antibody (clone HP6017, BioLegend) in 100 μL PBE and 30 minutes incubation with 1:20 Streptavidin MicroBeads (Miltenyi Biotec) in 100 μL PBE. Streptavidin MicroBeads captured yeast were positively selected by Multi-96 Columns (Miltenyi Biotec) placed in MultiMACS M96 Separator (Miltenyi Biotec). Selected yeast cells were recovered in 1 mL SDO−Ura at 30 °C for 24 hours.

#### NGS library preparation and sequencing

NGS sequencing was performed as previously described^20^. Briefly, DNA was extracted from yeast libraries using Zymoprep-96 Yeast Plasmid Miniprep kits (Zymo Research). Purified plasmids were amplified and indexed (2 rounds Phusion PCR, 24 cycles/round), pooled, gel purified, and sequenced on an Illumina NextSeq550 with 75 base pairs single-end sequencing. A minimum of 300k reads on each sample and 3 million reads for pre-selection library were collected.

### REAP data analysis

REAP score calculation was previous described^28^. A modified REAP scoring system was used in this study. The aggregate enrichment E_s_ (log fold change with zeroes in the place of negative fold changes) between the frequency of a protein in a sample and in the pre-selection library was calculated for each protein. Similarly, the aggregate enrichment E_b_ between the frequency of each protein in blank samples compared to the pre-selection library was also calculated. In order to account for background enrichment in the absence of sample, we subtracted blank enrichment E_b_ from sample enrichment E_s_. We defined REAP score as max(log[exp(E_s_)– exp(E_b_)],0) when E_s_ > E_b_, and 0 otherwise.

#### Autoantibody-treatment response association analysis

We assessed the association between the presence of each autoantibody identified in the cohort and the likelihood of patient having Schizophrenia. In instances with multiple replicates for a given sample, the mean REAP score was computed. To dichotomize the presence or absence of an autoantibody (AAb), REAP scores were binarized. A threshold of REAP = 1 was implemented, whereby: AAb = 1 indicates REAP 1 (presence) and AAb = 0 indicates REAP < 1 (absence).

For each autoantibody, Firth logistic regression was employed to calculate association with treatment response. The model was adjusted for known confounders age, sex, and medical comorbidities, represented as:

Schizophrenia status ∼ AAb + age + sex + race

Firth logistic regression, as opposed to conventional logistic regression, incorporates a penalty to the likelihood using the Jeffreys prior. This modification ameliorates challenges posed by low-frequency autoantibodies or situations of perfect separation. The regression analysis yielded adjusted odds ratios and associated p-values, computed using the logistf package in R.

#### Autoantibody discovery rate analysis

We performed a discovery rate analysis to assess the relationship between the number of patients in the cohort and the number of unique antigens with autoantibody reactivities observed. We took a random permutation of the patients. Beginning with the first patient in this permutation, we successively tallied the cumulative number of unique antigens exhibiting autoantibody reactivities as we incorporated data from each subsequent patient. For a more granular understanding, we maintained separate cumulative counts based on the frequency of antigen occurrence within the cohort, distinguishing common antigens (defined as antigens with a frequency greater than or equal to 0.01 in the cohort) from rare antigens (antigens with a frequency less than 0.01 in the cohort).

#### Autoantibody persistence analysis

We identified and tracked REAP hits exhibiting a REAP score greater than 1 on day 0. A REAP hit was defined to persist as long as it maintained a REAP score greater than or equal to 0.5. The persistence of a REAP hit concluded upon observing the first sample where the REAP score was less than 0.5 and no subsequent resurgence of the score above this threshold. In instances where REAP hits were observed in the last longitudinal sample of a patient without subsequent data to confirm persistence or decline, these hits were right-censored to account for potential truncation of data. To visually represent the persistence of REAP hits over time and estimate the median persistence duration, we employed the Kaplan-Meier survival curve. The curve, along with the median persistence time, was computed using the survival and survminer packages in R.

#### Lasso machine learning model

A Lasso model was trained to predict sample type (Schizophrenia versus healthy). The input for the model was binarized REAP reactivity scores from single samples for 971 healthy subjects and 352 Schizophrenia patients. To limit the number of features in the model, we only considered antigens observed in greater than 5%, resulting in 242 features. To assess model performance, we ran 10-fold cross-validation with hyperparameter tuning as part of the training pipeline. The performance of the models on each test fold was visualized using receiver operating characteristic (ROC) curves.

#### Tissue and cell-type contribution analysis

We quantified how much each tissue or cell type accounts for the disease-associated rise in autoantibody burden. We used the Human Protein Atlas (HPA) annotations to construct target sets for each tissue or cell type.

For each tissue or cell type, we constructed a set of X-enriched proteins from the HPA annotations: Group enriched, Tissue enhanced, or Tissue enriched. Per HPA, these terms are defined as:

- Tissue enriched: At least four-fold higher mRNA level in a particular tissue/cell type compared to any other tissues/cell types.
- Group enriched: At least four-fold higher average mRNA level in a group of 2-5 tissues or grouped cell types compared to any other tissues/cell types.
- Tissue enhanced: At least four-fold higher mRNA level in a particular tissue/cell type compared to the average level in all other tissues/cell types.

Proteins can belong to multiple tissues/cell types; by design, sets may overlap.

Subject-level burdens, for each subject :

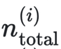: total number of reactive AAbs (all proteins).

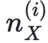: number of reactive AAbs whose target protein is in set (0 if none).

Define the between-group mean differences:

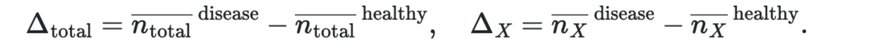

Percent contribution, for each:

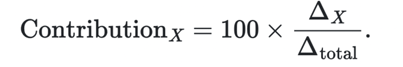

This reads as “the share of the overall AAb increase that is attributable to AAbs targeting.” Because HPA sets overlap, contributions across tissues/cell types need not sum to 100% and can exceed 100% or be negative when other sets decrease.

Uncertainty: we computed 95% confidence intervals via a stratified nonparametric bootstrap with replicates:

1. Resample subjects within status (healthy vs disease) with replacement to preserve group sizes.
2. Recompute the ratio for each resample.
3. The CI is the 2.5th–97.5th percentiles of the bootstrap distribution

### Autoantibody ELISA

250 ng of recombinant protein antigen in 100 μL PBS was added to wells of 96-well Immuno 2HB plates (Thermo Fisher Scientific). Plates were sealed and placed in 4°C overnight. After incubation, plates were washed once with 225 μL wash buffer (PBS + 0.05 % Tween 20). 150 μL blocking buffer (2 % HSA in PBS pH 7.0) was added to each well and incubated RT for 2 hours. Plates were then washed three times with 225 μL wash buffer. 100 μL plasma dilutions (starting with either 1:50 or 1:100 dilution, in blocking buffer) was added to the corresponding wells and incubated at RT for 2 hours. Plates were subsequently washed six times with 225 μL wash buffer. 1:5000 HRP anti-human IgG Fc (GenScript, #A00166) or isotype-specific antibody (Biolegend, 410603; Southern Biotech, IgG1: clone HP6001, IgG2: clone 31-7-4, IgG3: clone HP6050, IgG4: clone HP6025) in 100 μL blocking buffer was added to each well and incubated at RT for 1 hour. Plates were then washed six times with 225 μL wash buffer. 50 μL TMB substrate (BD Biosciences) 1:1 mixture was added to each well and developed at RT in the dark for 15 minutes. 50 μL 2N/1M H_2_SO_4_ was added to each well to terminate the TMB reaction. Absorbance at 450 and 570 nm was measured in a Synergy HTX Multi-Mode Microplate Reader (BioTek).

### Oxytocin Receptor Signaling assay

OXTR-expressing CHO-K1 cells (PathHunter® eXpress OXTR CHO-K1 β-Arrestin GPCR Assay) were thawed and plated following the manufacturer’s instructions and cultured overnight. As a positive control, OXTR-cells were cultured with varying concentrations of oxytocin ligand for 1.5 hours and read on a luminescence plate reader. The EC50 was determined and the concentration was used for subsequent antagonist assays. To assess antibody-mediated inhibition, cells were pre-incubated for 30 minutes with 91 µg/mL IgG purified from patient serum positive or negative for anti-OXTR autoantibodies. Media containing oxytocin ligand at the EC50 concentration was then added, and cells were incubated for an additional 1.5 hours. Cells were incubated with detection solution for 1 hour at room temperature in the dark and samples were read on a luminescence plate reader according to manufacturer instructions.

### 3D microfluidic blood–brain barrier (BBB) preparation

#### Model overview

The 3D human BBB model was established using OrganoPlate 3-lane-64 devices (MIMETAS) as published^39^. Each chip contained an extracellular matrix (ECM) gel in the central channel, a human brain endothelial microvascular tubule in the right channel, and a mixture of human primary astrocytes and pericytes seeded in the left channel, which migrated toward the endothelial compartment during culture on a bidirectional OrganoFlow rocker. Cells included in the model include:

- Human Brain Primary Endothelial Cells: Cell Systems, Cat: ACBRI 376, Lot: 376.03.02.01.2F
- Human Brain Vascular Pericytes: ScienCell, Cat: 1200, Lot: 33155
- Human Astrocytes: ScienCell, Cat: 1800, Lot: 33363

#### ECM preparation and endothelial channel coating

On day −1, the middle channel was filled with a collagen I/collagen IV mixture (4:1; each at 5 mg/mL), incubated at 37 °C for 15 minutes, and then overlaid with PBS to prevent desiccation. The right channel inlet was coated overnight with fibronectin (10 µg/mL) and collagen IV (10 µg/mL).

#### Endothelial cell seeding

On day 0, brain endothelial cells were dissociated and resuspended at 20,000 cells/µL. A total of 40,000 cells were introduced into the right channel outlet of each chip. Plates were placed at a 75° angle (manufacturer stand) and incubated at 37 °C for 6 hours to allow cells to attach along the ECM boundary. Outlets were then replenished with endothelial medium, and chips were transferred to the OrganoFlow rocker alternating between −7° and +7° every 8 minutes. Within 24–48 hours, brain endothelial cells formed an open tubular structure.

#### Astrocyte/pericyte channel coating and seeding

On day 1, the left inlet was coated with Matrigel. Primary astrocytes were resuspended in AM-PM medium (4:1 AM:PM, ScienCell) at 7,500 cells/µL, and primary pericytes were prepared similarly at 5,000 cells/µL. Cells were mixed at a 1:1 astrocyte:pericyte ratio, yielding 7,500 astrocytes and 5,000 pericytes per chip, and seeded through the left channel outlet. Plates rested for 4 hours at 37 °C before outlet wells were replenished with AM-PM medium and returned to rocker-based culture until day 10. Media was replaced every other day. Beginning on day 6, endothelial channels received standard EC medium lacking bFGF and VEGF.

#### Addition of patient / control sera

Patient or control sera diluted 1:20 in EC medium lacking bFGF and VEGF were added for 48 hours starting at day 9-10, followed by assessment of BBB function.

### 3D microfluidic BBB functional assays

#### Dextran and biocytin permeability

On day 10, 70-kDa dextran (0.5 µg/mL) and biocytin (0.5 µg/mL) were added to the right inlet and outlet wells (40 µL and 30 µL, respectively). Wells of the ECM inlet, left inlet, and left outlet were filled with 20 µL tracer-free medium. Plates were placed on the rocker and imaged every 15 minutes using the LICOR system. Fluorescence intensities were quantified on the Odyssey SA as published^39^. Permeability was expressed as normalized tracer intensity in the central ECM well:

Normalized intensity = (Sample ECM middle-well intensity / Mean ECM-only control middle-well intensity) × 100

ECM-only chips served as controls for passive leakage.

#### TEER measurements

TEER was assessed on day 10 using the OrganoTEER device, applying the manufacturer’s low-TEER setting optimized for endothelial barriers as published^39^.

### Cav3.3 electrophysiology model

#### Cav3.3 cell line

HEK293 cells stably expressing hCav3.3 channels were generously provided by Jen Q. Pan. Expression of hCav3.3 in these cells is controlled by a tetracycline-responsive promoter, and can be induced by doxycycline. Three days prior to recordings, cells were split with fresh complete media with 2μg/mL doxycycline, which was refreshed 48 hours before recordings. The complete media contained: DMEM (Cytiva #SH30284.01), 10% fetal bovine serum (Corning #35-016-CV), 1x GlutaMAX (Gibco #35050061), 1x MEM Non-Essential Amino Acids Solution (Gibco #11-140-050), and 1x Penicillin-Streptomycin (Gibco #15140122).

#### IgG 24-hour incubation

At 24 hours before recordings, the hCav3.3 HEK293 cells were split and plated at a density of ∼20,000/well in a 24-well plate with 0.1% gelatin solution (Millipore) treated glass coverslips in each well. The media also contained 2μg/mL doxycycline and IgG from healthy donors or schizophrenic patients REAP+ for anti-Cav3.3 autoantibodies. The final concentration of IgG was between 100-150μg/mL. Cells were maintained in an CellXpert® incubator (Eppendorf) at 37°C and 5% CO2 until recording. Coverslips containing the cells were moved to a chamber immediately prior to recordings.

#### Cav3.3 Electrophysiology

Calcium currents were recorded whole cell in the voltage clamp configuration using HEK293 cells stably expressing hCav3.3 channels. Cells were patched using borosilicate glass pipettes with resistances of 2-5MΩ made using a P-1000 micropipette puller (Sutter Instrument). Recording pipettes were filled with an internal solution that contained the following (in mM): 110 CsF, 10 CsCl, 10 NaCl, 10 EGTA, and 10 HEPES at a pH of 7.2 (Nanion Technologies). The external recording solution contained the following (in mM): 150 NaCl, 10 HEPES, 2.5 KCl, 1.25 NaH2PO4, 2 CaCl2, 1 MgCl2, and 5 Glucose, adjusted to a pH of 7.25 with NaOH. HEK293 cells were visualized under differential interference contrast using a BX51WI microscope and a 40X objective (Evident Scientific). Only cells not in direct contact with nearby cells were chosen for recording. Prior to break-in, the holding potential was adjusted to −90mV and a gigaseal was obtained. Series resistance was compensated to 70% (prediction and correction). Channel activation was elicited by applying 100ms voltage steps from −90mV to +20mV in increments of 10mV. P/-4 leak subtraction was applied to all currents online. All recordings were performed at room temperature. Data were collected using pClamp11.3 software (Molecular Devices), a Multiclamp 700B amplifier (Molecular Devices), and a Digidata 1550B digitizer (Molecular Devices). Currents were recorded at a sampling rate of 20kHz and low-pass filtered at 1kHz prior to analysis.

#### Data analysis

Data were analyzed using Clampfit 11.3 (Molecular Devices), Excel (Microsoft), and custom written Matlab code (MathWorks). Peak currents were measured as the most negative current value within each voltage step and current densities were calculated by dividing the peak current at each voltage step by the measured cell membrane capacitance.

### Statistics and reproducibility

Statistical analyses were conducted using R (v4.2.3) and GraphPad Prism (v10.0.0) and are described in the figure legends and Methods. No statistical methods were used to predetermine sample size. The REAP assay was performed in duplicate with two technical replicates for all samples on separate days as previously described^28^. All in vitro ELISA and signalling assays were conducted in duplicate with two technical replicates The number of repetitions for other experiments is provided in the figure legends.

## DATA AVAILABILITY

All data used to generate figures and tables in this study are included in the Source Data. REAP data will be made available on reasonable request from the corresponding authors, subject to restrictions related to patient privacy in accordance with institutional policies and the Health Insurance Portability and Accountability Act. Source data will be provided with this paper prior to publication.

## CODE AVAILABILITY

The custom code for the analysis of REAP data will be made available on GitHub prior to publication.

## ACKNOWLEDGEMENTS

We thank all members of the Ring laboratory and Seranova for technical assistance and helpful discussions. A.M.R. was supported by grants from the Mark Foundation for Cancer Research and the Pew Charitable Trusts, and gifts from the Anderson and Bezos Families. J.R.J. was supported by the Yale Medical Scientist Training Program. U.A. and D.A. were supported for this work by the following funding: NHLBI R61/R33 HL159949, NEI R01EY033994, PANDAS Network, PANDAS Physician Network and NIH NCAT TL1TR001875 (UA).

## AUTHOR CONTRIBUTIONS

K.N., J.R.J., D.C.G., and A.M.R. conceived the study and designed the research. K.N., D.C.G., and A.C.L. provided clinical resources and patient samples. J.R.J., S.M.Y., K.Q., A.A.N., S.M.C., U.A., B.M.S., and N.H. carried out experimental studies. J.R.J. and L.F., performed data analysis. B.S.M., R.W.T., and D.A. contributed supervision and expertise for specialized BBB studies and edited the manuscript. D.C.G. and A.M.R. supervised the work. J.R.J., K.N., D.C.G., and A.M.R. drafted the manuscript, with input from all authors. All authors reviewed and approved the final version.

## COMPETING INTERESTS

A.M.R. is an inventor of a patent application assigned to Yale University describing the REAP technology (patent number: WO2021189053A1). In addition, A.M.R. is the founder and a director of Seranova Bio, the commercial licensee of REAP. S.M.C., L.F., B.S.M. are employees of Seranova Bio. J.R.J. K.N., U.A., D.A., S.M.Y., B.M.S., N.H., R.W.T and D.C.G.. declare no competing interests.

## SUPPLEMENTARY DATA

**Supplementary Figure 1:**
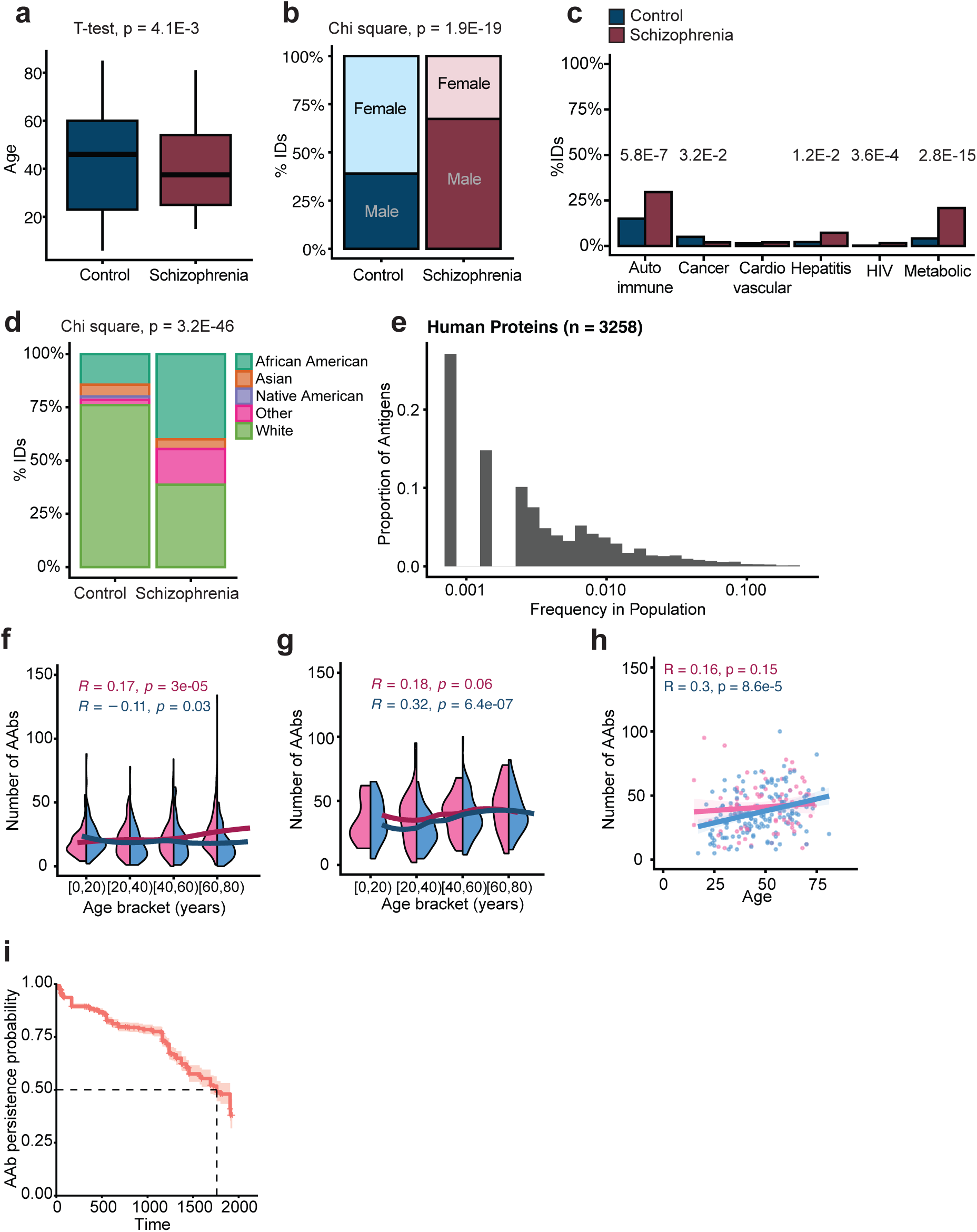
**A.** Age of individuals by group (control, n=971; schizophrenia, n = 352). Significance was assessed using unpaired two-sided t test. The central lines indicate the group median values, the top and bottom lines indicate the 75th and 25th percentiles, respectively, the whiskers represent 1.5× the interquartile range. **B.** Sex of individuals by group (control, n=971; schizophrenia, n = 352). Significance was assessed by Chi Square test. **C.** Percentage of individuals from control (n=971) and schizophrenia (n=264) cohorts with listed comorbidities. Significance assessed by fisher’s exact test and adjusted for multiple comparisons by Benjamini-Hochberg. HIV = Human immunodeficiency virus. **D.** Percentage of individuals from control (n = 971) and schizophrenia (n=352) belonging to listed race. Significance assessed by Chi square test. **E.** Autoantibody frequency distribution across the cohort. Each bar represents the proportion of autoantibodies observed at the specified frequency within the cohort. **F, G.** paired violin plot depicting number of autoantibodies (AAbs) per individual by age bracket within the control cohort **(F)** and schizophrenia cohort **(G).** Pink = female; blue = male. Moving average depicted by magenta and blue lines for female and male, respectively. Correlation (pink = female, blue = male) was assessed using Spearman’s correlation **H.** The relationship between number of autoantibody (AAb) reactivities per individual and age, grouped by sex, among the chronic psychosis population (n = 247). Correlation was assessed using Spearman’s correlation. The magenta and blue lines show the linear regression for females and males, respectively, and the shading shows the 95% CIs. Each dot represents one individual (pink = female, blue = male). **I.** Kaplan–Meier survival curve representing the persistence of autoantibodies (AAbs) (n = 5,701 reactivities). Persistence was defined as maintaining a REAP score >= 0.5 in the longitudinal samples for reactivities detected at the first time point for a given subject with REAP score >= 1. Data are presented as survival probability with 95% confidence intervals. Dashed lines indicate the time points after treatment at which 50% of the AAb reactivity was still detectable.

**Supplementary Figure 2:**
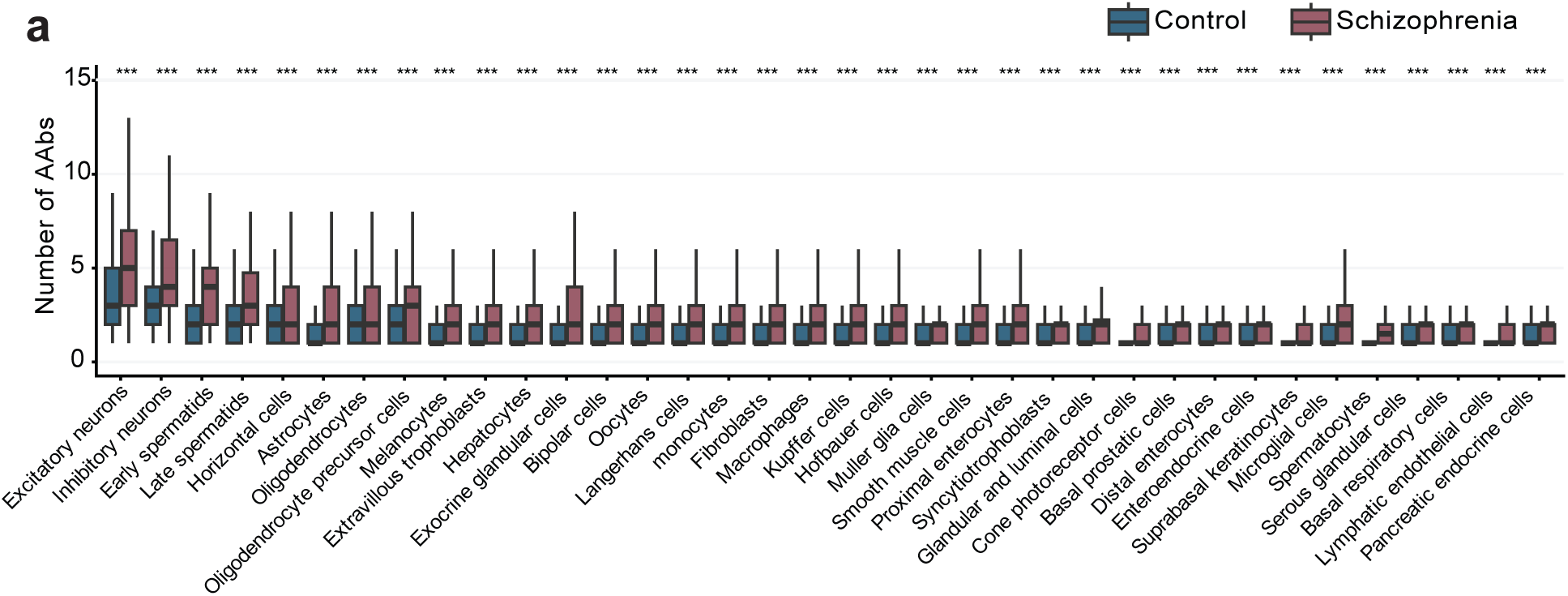
**A.** The number of autoantibody (AAb) reactivities against each tissue category per individual by group (control, n = 971; schizophrenia = 352). Single cell categories are composed of REAP reactivities bucketed by human protein atlas mRNA expression data. Significance was assessed by unpaired two-sided Wilcoxon with correction for multiple hypotheses by Benjamini-Hochberg. *** ∼ <0.001, ** ∼ <0.01, * ∼ <0.05.

**Supplementary Figure 3:**
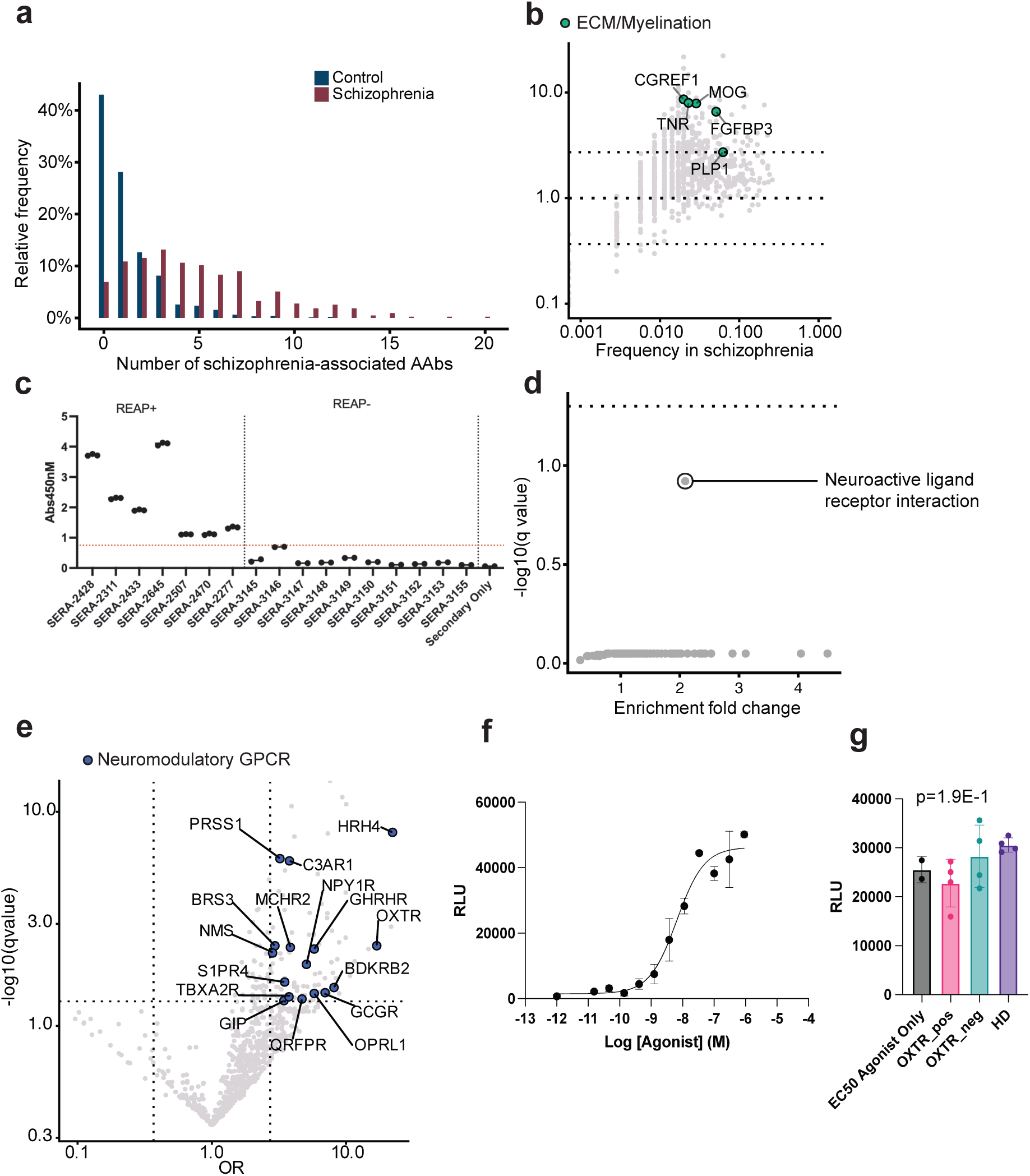
**A.** requency of individuals from control (n=971) and schizophrenia (n=352) cohorts with the listed number of schizophrenia-associated autoantibodies (AAbs). Schizophrenia-associated autoantibodies are defined as estimate >1 (OR >2.72) and q-value <0.05. **B**. Frequency plots depicting all autoantibody reactivities detected, with proteins belonging to the ECM/myelination group from figure 3B bolded and colored. Horizontal dashed lines indicate OR threshold of +/− 2.72. Each dot represents one reactivity. **C.** ELISA validation of binding for the 3rd extracellular domain HCN1. Each dot represents one technical replicate. Each column represents one patient. REAP+ = HCN1_Epitope_3 REAP score >= 1; REAP-= HCN1_Epitope_3 REAP score <1. **D.** KEGG pathway analysis of 93 autoantibodies enriched in schizophrenia against the REAP background library. Each dot represents one pathway. Dashed line represents significance threshold after correction for multiple comparisons by FDR. **E.** Volcano plot depicting all autoantibody reactivities detected, with reactivities that are significantly enriched in schizophrenia (q-value <0.05, odds ratio (OR) >2.72) and target proteins belonging to the neuroactive ligand receptor interaction KEGG pathway bolded and colored. Each dot represents one autoantibody reactivity. Vertical dashed lines indicate OR threshold +/−2.72. Horizontal dashed line indicates q-value threshold of 0.05. **F.** titration curve for OXTR beta-arrestin chemiluminesince assay. Each point represents 2 measurements. **G**. chemiluminescence signal (RLU) for OXTR assay at 91 ug/ml OXTR ligand. Each dot represents plasma from one individual and an average of 2 technical replicates. Significance was assessed by one way ANOVA. OXTR_pos = OXTR AAb positive schizophrenia patient; OXTR_neg = OXTR AAb negative schizophrenia patient.

**Supplementary Figure 4:**
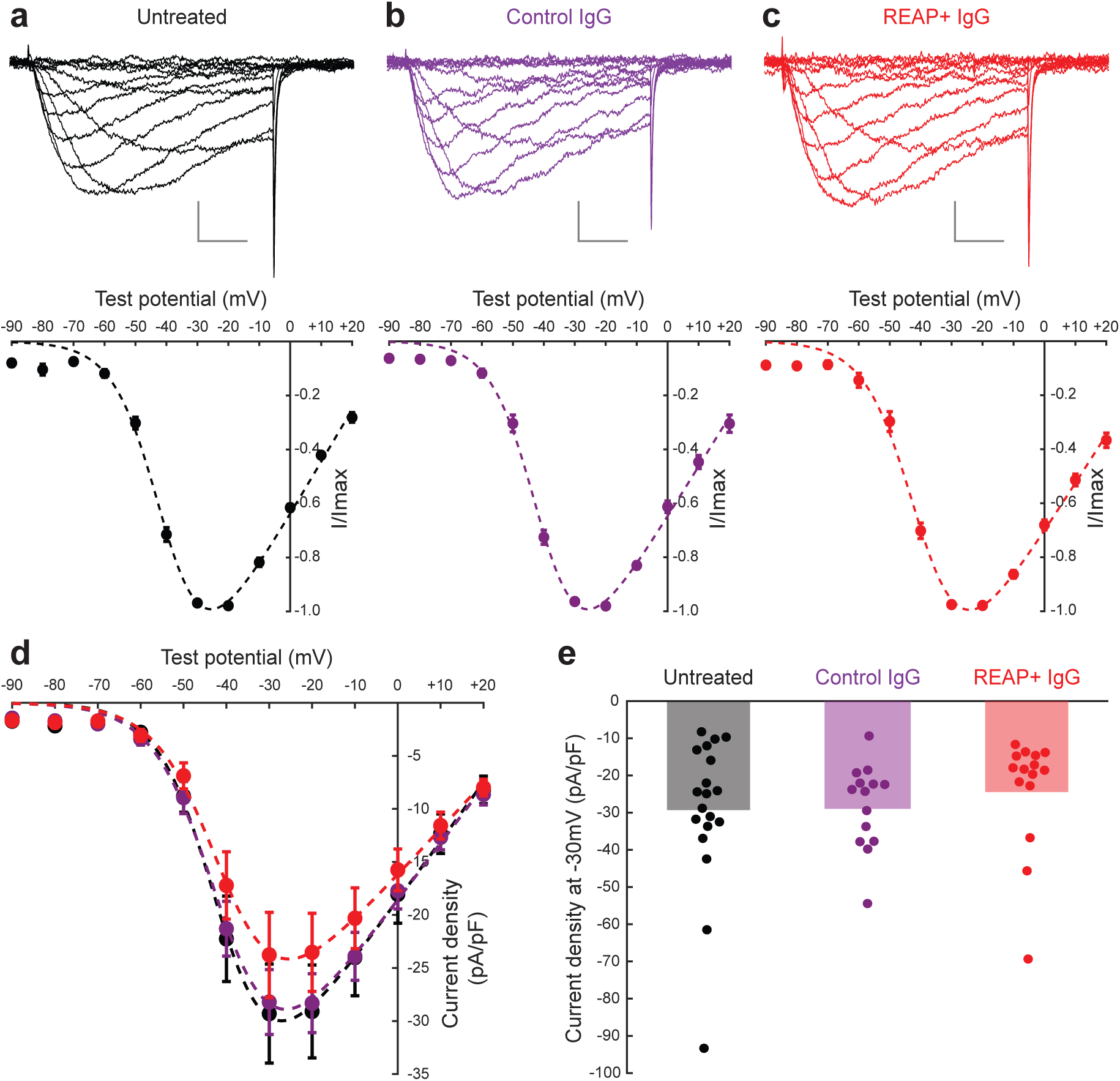
Cav3.3 currents recorded from HEK293 cells following 24-hour incubation with no IgG, IgG from healthy controls, or IgG from schizophrenic patients REAP+ against Cav3.3. **A.** (Top) Example Cav3.3 mediated currents from one HEK293 cell not treated with IgG. Currents were elicited by 100ms voltage steps from −90mV to +20mV in increments of 10mV. (Bottom) IV curve for Cav3.3 mediated currents from all recorded cells not treated with IgG. Currents from each cell were first normalized and then averaged; bars indicate SEM. The dotted line indicates a fit to I_norm_ = (G_max_)*(V_m_-V_rev_)/[1+exp(V_1/2_-V_m_)/k]],^49^ where G represents conductance, V_m_ represents the step potential, V_rev_ represents the apparent reversal potential, V_1/2_ represents activation midpoint, and k represents a slope factor. **B.** Same as A, but showing data from cells incubated for 24 hours with IgG from healthy controls. **C.** Same as A, but showing data from cells incubated 24 hours with IgG from schizophrenic patients and REAP+ against Cav3.3. For A-C, the vertical scale bar indicates 10pA/pF and the horizontal scale bar indicates 20ms. **D.** IV curve showing the relationship between current density (pA/pF) and test potential, where the average values for all cells within a treatment group are plotted and fit as in A-C; bars indicate SEM. Data from untreated, control IgG treated, and patient REAP+ IgG treated are shown in black, purple, and red, respectively. **E.** Peak current density values from each treatment group at a test potential of −30mV. Dots indicate data from individual cells and bars indicate average values. Significance was assessed by Kruskal Wallis (p=0.233). For all data shown, n=19 for untreated cells, n=14 for control IgG treated cells (from 5 donor samples), and n=15 for REAP+ IgG treated cells (from 4 patient samples).

**Supplementary Figure 5 A, B, C.**
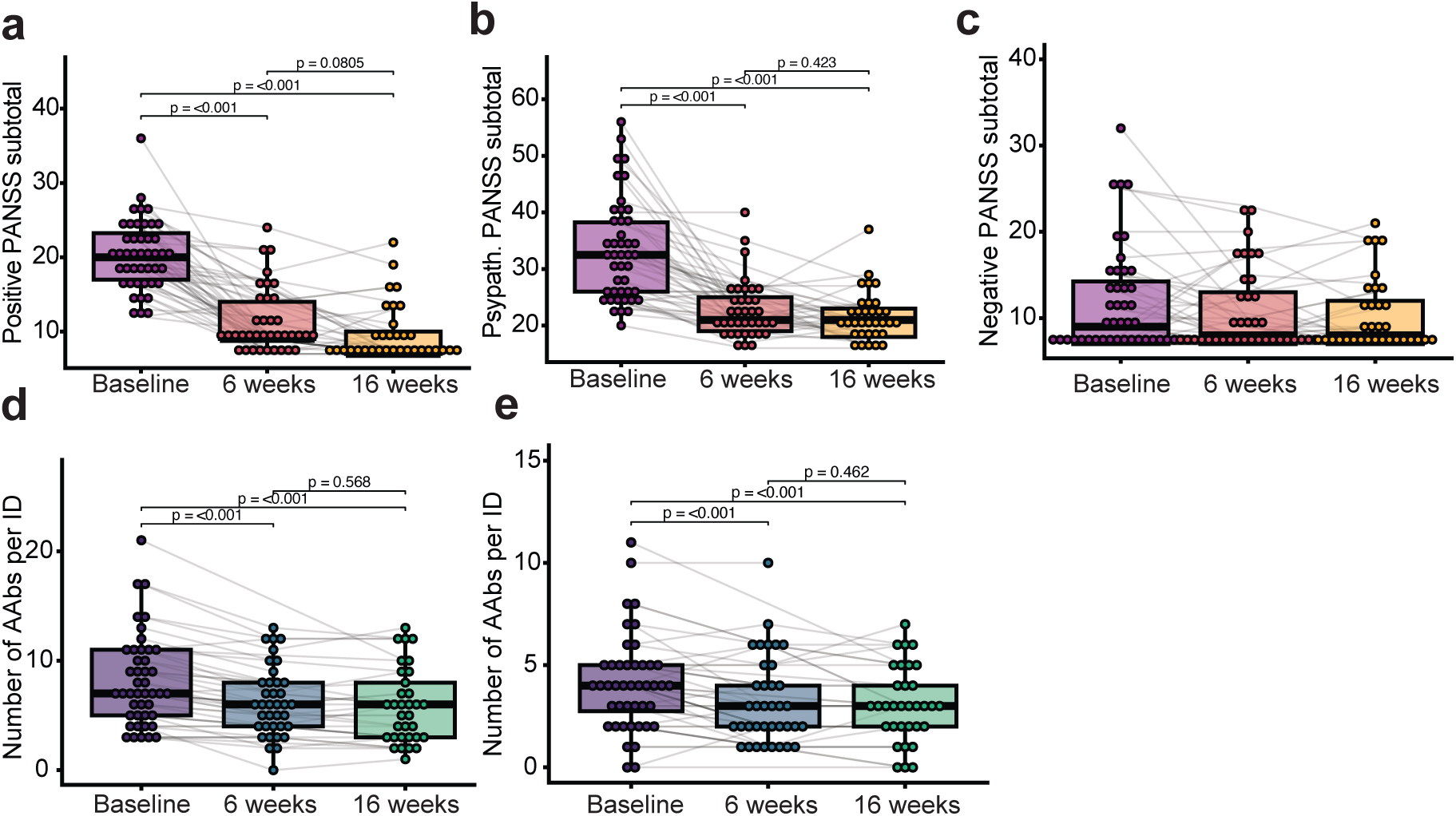
PANSS subscore for positive **(A)**, general psychopathology **(B)**, and negative **(C)** domains for participants (n = 44) during the study. Data were modeled using a linear mixed-effects model (n=44 patients) with timepoint as a fixed effect and subject identity as a random intercept. The overall effect of time was assessed via a Type III Wald F-test with Kenward-Roger degrees of freedom approximation (A: F(2, 74) = 955.5, p = <2.2E-16), (B: F(2, 74.9) = 45.0, p = 1.4E-13), (C: F(2, 72.3) = 1.4, p = 0.25). Pairwise comparisons were performed using estimated marginal means with p-values adjusted for multiple comparisons using the Holm-Bonferroni method. Each dot represents one individual per time point, with multiple time points for a given individual connected by grey lines For the box plots, the central lines indicate the group median values, the top and bottom lines indicate the 75th and 25th percentiles, respectively, the whiskers represent 1.5× the interquartile range. **D, E**. Number of brain- and excitatory or inhibitory neuron-enriched (**D**) and schizophrenia-associated (**E**) autoantibody (AAb) reactivities per individual (ID) at baseline, 6 weeks, and 16 weeks of risperidone treatment. Data were modeled using a linear mixed-effects model (n=44 patients) with timepoint as a fixed effect and subject identity as a random intercept. The overall effect of time was assessed via a Type III Wald F-test with Kenward-Roger degrees of freedom approximation (D: F(2, 68.7) = 33.5, p = 7.1E-11), (E: F(2, 69.3) = 12.2, p = 2.8E-5. Pairwise comparisons were performed using estimated marginal means with p-values adjusted for multiple comparisons using the Holm-Bonferroni method. Each dot represents one individual per time point, with multiple time points for a given individual connected by grey lines. For the box plots, the central lines indicate the group median values, the top and bottom lines indicate the 75th and 25th percentiles, respectively, the whiskers represent 1.5× the interquartile range.

